# Plasmodesmata-like intercellular connections by plant remorin in animal cells

**DOI:** 10.1101/791137

**Authors:** Zhuang Wei, Shutang Tan, Tao Liu, Yuan Wu, Ji-Gang Lei, ZhengJun Chen, Jiří Friml, Hong-Wei Xue, Kan Liao

## Abstract

Plasmodesmata (PD) are crucial structures for intercellular communication in multicellular plants with remorins being their crucial plant-specific structural and functional constituents. The PD biogenesis is an intriguing but poorly understood process. By expressing an *Arabidopsis* remorin protein in mammalian cells, we have reconstituted a PD-like filamentous structure, termed remorin filament (RF), connecting neighboring cells physically and physiologically. Notably, RFs are capable of transporting macromolecules intercellularly, in a way similar to plant PD. With further super-resolution microscopic analysis and biochemical characterization, we found that RFs are also composed of actin filaments, forming the core skeleton structure, aligned with the remorin protein. This unique heterologous filamentous structure might explain the molecular mechanism for remorin function as well as PD construction. Furthermore, remorin protein exhibits a specific distribution manner in the plasma membrane in mammalian cells, representing a lipid nanodomain, depending on its lipid modification status. Our studies not only provide crucial insights into the mechanism of PD biogenesis, but also uncovers unsuspected fundamental mechanistic and evolutionary links between intercellular communication systems of plants and animals.

**Significance:** Remorin is rising as a crucial lipid microdomain marker, as well as an essential component of plasmodesmata in plants. However, the biological role of remorin in plants is elusive. With a heterologous system, we found that remorin expression is able to form intercellular filamentous structure, namely remorin filament (RF), connecting neighboring mammalian cells functionally. By employing multiple approaches, we tested the functionality of RFs, as well as investigated their structures. RFs highly resemble plant plasmodesmata, in many ways, such as its morphology, molecular constitutes, and its ability to transport macromolecules intercellularly. Our study provides novel insights into the biogenesis of plasmodesmata and uncovers fundamental evolutionary links in molecular construction of intercellular connections in both plants and animals.

## Introduction

In a multicellular organism, cells need to exchange both substances (such as proteins, sugars and nuclear acids) and information (such as hormones and secondary messengers) to coordinate with each other for normal cell activity and coordinated tissue development. Cell-to-cell communication is essential in both plants and animals; however, the involved subcellular structures and underlying molecular machineries are fairly diverse (1, 2), presumably owing to the fact of independent evolution of multicellularity in these two major eukaryotic groups and very different cellular structures and tissue morphology. The vesicle systems of plants and animals transport biomolecules/materials to neighboring cells through phagocytosis, pinocytosis, or receptor-mediated endocytosis. A complex system specific for plants, plasmodesmata (PD), executes more efficient intercellular communication, enabling plants to have a tightly controlled symplasmic network crossing the apoplast, whereas animal cells without complication of cell walls can rely on direct membrane connections.

PD are nanotubes connecting neighboring cells in plants and given their absence in animals it is generally considered to be one of the major differences between animal and plant tissues (1–3). Previous studies showed that remorin is a plant-specific protein located at the PD that marks the lipid microdomain or nanodomain, it has also been reported to be related to the function of plant PD (4–9). In rice (*Oryza sativa*), *GRAIN SETTING DEFECT1* (*GSD1*), encoding a remorin protein, regulates PD conductance to participate in the grain setting process (5). GSD1 functions through interacting with OsACTIN1 and OsPDCB1 (PLASMODESMATA CALLOSE BINDING PROTEIN1) (5). A recent study reveals that *Arabidopsis* remorin proteins, REM1.2 and REM1.3, are crucial regulators for plasma membrane nanodomain assembly to control PD closure, in response to a defense hormone salicylic acid (SA) (9). Based on these observations, remorin might be a key factor for PD biogenesis. However, due to the essential physiological function of the plant PD system, it is difficult to verify the molecular role of remorin in PD function with the plant system.

Here, we investigated the effects of heterologous expression of remorin in mammalian cells. This leads to the formation of PD-like structures connecting cells and allowing exchange of material. Our study shows that a single plant protein, remorin, is sufficient to induce functional PD-like intercellular connections in animal cells providing crucial insights not only into the mechanism of PD biogenesis but also into the evolutionary commonalities in intercellular communication between animals and plants.

## Results

### Deficiency of *REMORINs* leads to PD morphological abnormality in *Arabidopsis*

In *Arabidopsis*, the remorin family comprises 16 members, of which Remorin1.2 (REM1.2, AT3G61260) and Remorin1.3 (REM1.3, AT2G45820) show high similarities and were reported to be localized at the PD (4, 6, 7, 9). To study the role of remorin in PD formation and function, we obtained a T-DNA insertional mutant for *rem1.3* and a CRISPR-CAS9 mutant for *rem1.2*, and generated also a double mutant *rem1.2 rem1.3* (Fig. S1A, B). Notably, the *rem1.2 rem1.3* double mutant exhibited attenuated growth (Fig. S1C), indicating that REM1.2 and REM1.3 are essential for plant growth. This is in line with the previous notion that the *rem1.2 rem1.3* double mutant also showed attenuated growth at the seedling stage (9). We firstly tested the potential involvement of PD in the remorin function. PD play an essential role in transporting different molecules, including carbohydrates in plant cells (1, 2). Notably, 51% of the leaves of the *rem1.2 rem1.3* had a significantly positive starch accumulation whereas undetectable in WT (Fig. S1D), indicating its deficiency in starch unloading. A further test for the PD permeability with a fluorescent dye (Fig. S1E), FDA (fluorescein diacetate) (5, 10) confirmed that, FDA was pre-infiltrated into plants and then bleached by a strong laser exposure. Intriguingly, the *rem1.2 rem1.3* plants recovered more slowly than WT (WT, 70%; *rem1.2 rem1.3*, 30% at 290 s).

Furthermore, detailed analysis with TEM (transmission electron microscopy) indicated that the number of PD was significantly decreased in *remorin* mutants (Fig. S1F; WT, 5.4; *rem1.3- 1*, 4.0; *rem1.2-1*, 2.1; *rem1.2 rem1.3*, 1.8). However, several dark osmiophilic globules (∼ 50 nm in size) were observed in the *rem1.3* and *rem1.2* cells, with a greater number in the *rem1.2 rem1.3* cells (Fig. S1F), which is probably due to PD dysfunction (9, 11). In consistent with the notion that REM1.2 and REM1.3 also play a crucial role in the closure of PD (5, 9), all these defects support the essential function of remorin in PD.

### Filamentous intercellular structures by expressing remorin in mammalian cells

In line with the notion that remorin localizes to PD and regulates PD aperture (9), our observations support that the requirement of remorin for PD function. To gain additional insights into remorin function, we transfected eGFP-tagged *REM1.3* into the mammalian cells. We hypothesize that this heterologous system might be helpful to understand the exact cellular function of remorin, for the relatively less complicated background and the absence of all other plant-specific factors.

Introducing eGFP-REM1.3 into Cos7 cells, confocal laser scanning microscopy (CLSM) observation revealed eGFP-REM1.3 distribution in some filamentous structures (termed remorin filaments, RFs) connecting neighboring cells (Fig 1A). Interestingly, both ends of the RFs were swelling, with positive staining of the endoplasmic reticulum (ER) marker protein calnexin (12) accumulating at both ends. This became particularly apparent when the cell density is low. A schematic diagram of RFs across two animal cells (Fig 1A right) is a typical structural feature for plant PD that also included ER structures more prominent at the endos (9). Therefore, we hypothesize that these RFs possibly assemble into structures similar to plant PD. To directly observe their biogenesis, a time-lapse imaging was obtained using a video microscope. The results clearly show how the RFs bud out of the vesicles and grow by gradual elongation (Fig 1B).

**Fig 1.**
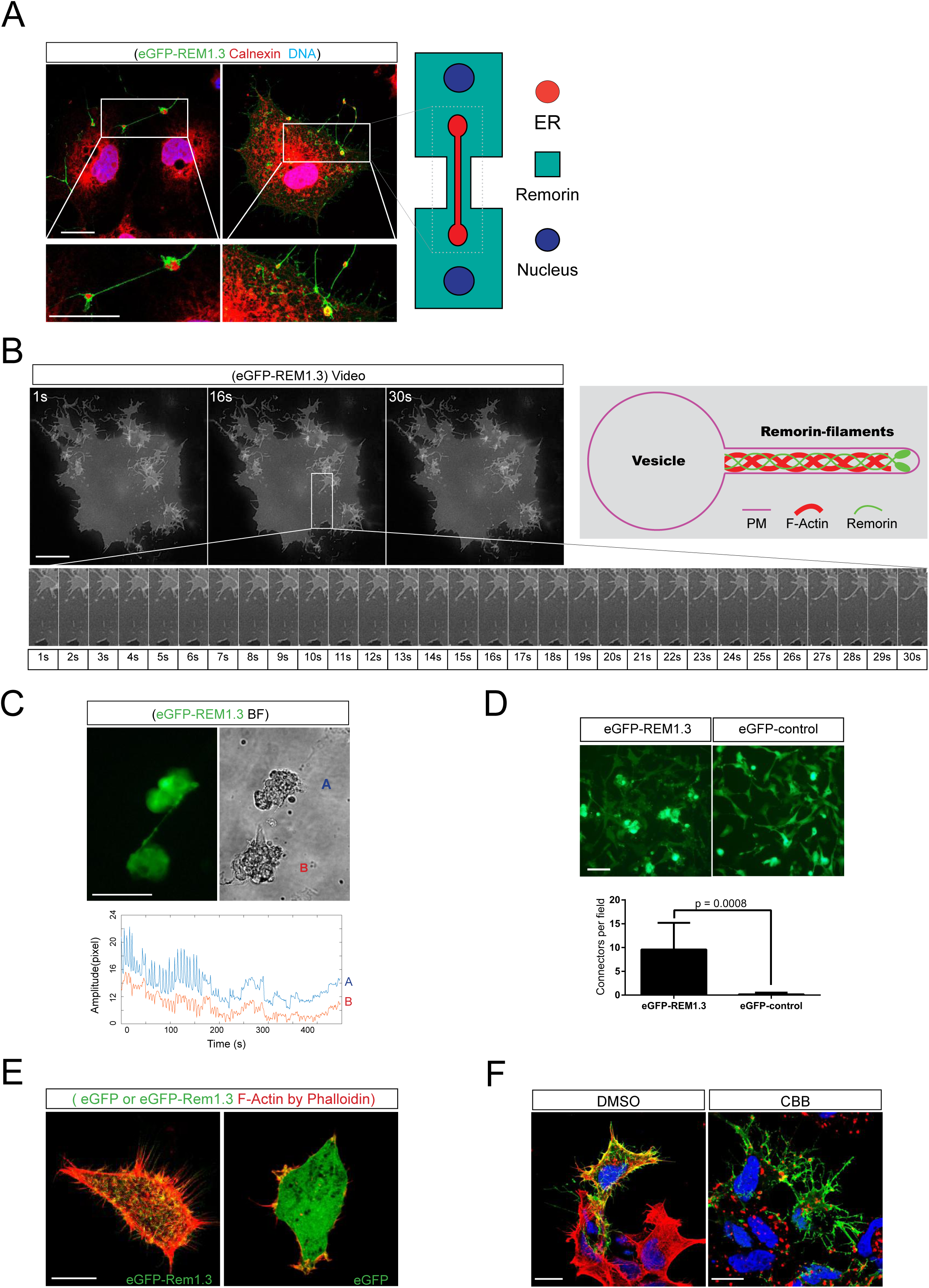
PD-like structure formed in animal cells transfected with eGFP-REM1.3. (A) PD-like structure formed in animal Cos7 cells transfected with eGFP-REM1.3 and stained for ER with calnexin antibody (red: Calnexin; green: REM1.3; blue: DNA; scale bar, 10 μm). The right portion shows PD-like structure pattern on animal cells (red: endoplasmic reticulum; green: cell body; blue: cell nucleus). (B) Image subtraction of GFP-REM1.3-transfected Cos7 cells by video microscope (gray: REM1.3; scale bar, 10 μm). The upper right panel shows the pattern diagram of remorin budding out the vesicle. (C) Mouse cardiac myocytes infected with eGFP-REM1.3 or eGFP control vector adenovirus. Connections are two or more cells or cell groups linked by filamentous structures with collaborative impulse. The pulsatile displacement of both cell clusters (red: A; blue: B) was recorded on the red and blue lines on Image Pro Plus software, respectively. (green: REM1.3 or eGFP-control; gray: bright field; scale bar, 200 μm). (D) Statistical analysis of cell populations or cells that are connected together (Numbers of connectors per field ± SD). (E) HEK293 cells transfected with eGFP-REM1.3 (green: REM1.3; red: phalloidin-stained F-actin; scale bar, 10 μm). (F) F-actin depolymerization prompted collapse of RFs. eGFP-REM1.3-transfected HEK-293 cells were treated with the depolymerization reagent cytochalasin B (1 μg/mL) for 15 min (green: REM1.3; red: phalloidin-stained F-actin; blue: DNA; DMSO: solvent control; CBB: cytochalasin B, scale bar, 10 μm).

The main function of PD in plants is to communicate through signals and biomolecules between cells (1–3, 13). To confirm that remorin expression in animal cells can create functional cell-to-cell connections, we infected primary myocardial cells, using the eGFP-REM1.3 adenovirus. Because primary myocardial cells have a rhythmic impulse, it is possible to demonstrate that two cells or whole cell populations are connected if they have coordinated rhythm. In line with this, simultaneous video analysis showed that two cells or cell populations linked by RFs share a synchronous beat rhythm revealing that ions pass through the two groups of cardiomyocytes through the RFs consistent with the functional intercellular connections compared to the control (free eGFP) cells (Fig 1C and D).

These experiments reveled that expression of plant remorin in different types of animal cells produce RF cell-to-cell connections similar to plant PD.

### Parallels between remorin-induced connections in animal cells and plasmodesmata

To characterize the similarity between RFs and the plant PD and to explore their biogenesis, we performed a series of microscopic observations and biochemical experiments. RFs have a diameter of approximately 100-150 nm, one-tenth to half of a cell length, and are usually branched (Fig 1E, 4A-B and Table S1). Co-stained with different markers, RFs stained positive for phalloidin (actin filaments, AFs) and negative for α-tubulin (microtubule, MT) markers, vimentin (intermediate filament) and Fascin (filopodial markers), suggesting that RFs are AF-based (Fig 1E). Cells treated with cytochalasin B, an AF-depolymerizing reagent, exhibited collapsed RFs (Fig 1F), confirming that RFs not only contain AFs and also that AFs are essential for RF integrity. These characteristics are very similar to plant PD, which was also found to be associated with enriched actin filaments (1, 3, 14–16). Indeed, these observations may also help elucidate the role of actin filaments in PD function, here via interacting with the remorin protein.

To further explore the analogy between the RFs in animal cells and PD in plants, we analyzed the basic biochemical properties of remorin in animal cells. Remorin was previously reported to localize at certain lipid microdomains (4, 5, 7). In consistent with the previous results, remorin isoforms (REM1.2, REM1.4, and REM1.1) can be expressed in mammalian cells and is associated with the lipid microdomains through sucrose-density gradient centrifugation (Fig S2A). Remorin is also reported to be post-translationally modified by palmitoylation (7). We employed an acyl-biotin exchange assay to detect the palmitoylation of REM1.3. The results showed that wild-type REM1.3 expressed in mammalian cells can be palmitoylated (Fig S2B). These results showed that heterologously expressed REM1.3 in mammalian cells presents the same subcellular localization and biochemical modification as in plant cells.

These experiments reveled that plant remorin-induced RF cell-to-cell connections in animal cells are actin-based like PSs and are structurally and biochemically similar to plant PD.

### Remorin-induced functional intercellular connections in mammalian cells

To test whether heterologous expression of remorin in animal Cos7 cells can result in communication/transport of macromolecules between cells to mimic the PD function. A set of highly sensitive detection systems based on Cre-LoxP was designed (Fig 2A). The lmlg (LoxP-mCherry-multiple stop codons-LoxP) element is an expression box consisting of LoxP, mCherry, multiple stop codon sites, LoxP and GFP proteins in series. In the absence of Cre recombinase, only mCherry was expressed (red); in the presence of Cre recombinase, the mCherry and multi-stop codon sites in the middle of LoxP were excised, leading to expression of the GFP protein (green). The Cre recombinase was transfected into donor cells. If the proteins between the donor and recipient cells could communicate with each other, receptor cells with green fluorescence were expected to be visualized.

**Fig 2.**
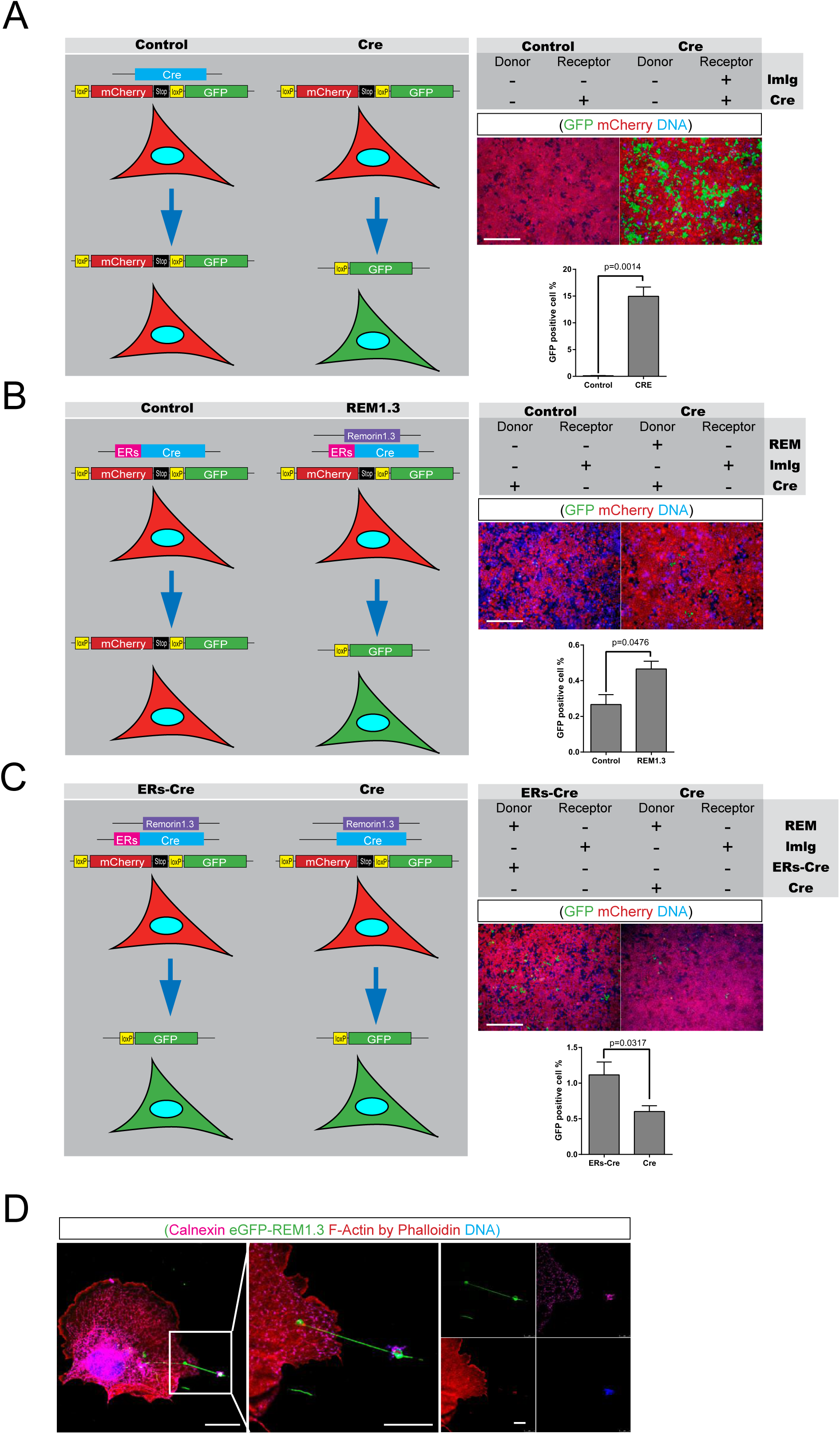
Exogenously expressed remorin enables animal cell-cell material transport. (A-C) Co-expression of remorin and endoplasmic-reticulum-located CRE recombinase (ERs-Cre) in donor cells, expression of “lmlg” (LoxP-mCherry-multiple stop codons-LoxP) elements in acceptor cells. The lmlg element is an expression box consisting of LoxP, mCherry, multiple stop codon sites, LoxP and GFP proteins in series. In the absence of Cre recombinase, only mCherry was expressed (red); in the presence of Cre recombinase, the mCherry and multi-stop codon sites in the middle of LoxP were excised, leading to expression of the GFP protein (green). The Cre recombinase was transfected into donor cells. If the proteins between the donor and recipient cells could communicate with/transport to each other, the lmlg element is cleaved by CRE, mCherry (mCr) and multiple stop codons are excised, and GFP begins to express, receptor cells with green fluorescence were expected to be visualized. After transfection for 48 hours, 1:1 mixed cells are cultured for 48 hours before observation by microscope (green: GFP; red: mCherry; blue: DNA: Scale bar, 100 μm), double-positive cell statistics conducted by flow cytometry with a bar chart [% Numbers of GFP positive / (GFP positive + mCherry positive) cells ± SD]. (D) PD-like structure transports DNA from cell. Animal cells transfected with eGFP-REM1.3 (green: REM1.3; red: phalloidin stained F-Actin; pink: Calnexin; blue: DNA stained by diaminophenindole, DAPI; scale bar, 10 μm)

Analysis by fluorescence microscopy and flow cytometry showed that donor cells expressing remorin and an ER-localized Cre recombinase enabled the receptor cells to exhibit GFP fluorescence (Fig 2B and C), while non-ER-localized Cre recombinase or a lack of remorin expression in the donor cells exhibited no detectable signal (Fig 2B and C). This result confirms that remorin expression in the donor cells can transmit Cre recombinase to the receptor cell (Fig 2B and C). Next, by DNA staining, we observed that RFs were able to transport DNA between cells (Fig 2D).

These observations collectively show that RFs indeed represent functional intercellular connections capable of transmitting not only electrical signals, but also proteins and DNA between animal cells.

### Role of different remorin domains in mammalian cells and plants

To study the biogenesis process of RFs, we investigated how the primary structure of REM1.3 contributes to RF formation in mammalian cells. We expressed various truncated REM1.3s (Fig 3A and S5), and observed RF formation and lipid microdomain localization to gain insight into the contribution of each domain as summarized in Table S2.

**Fig 3.**
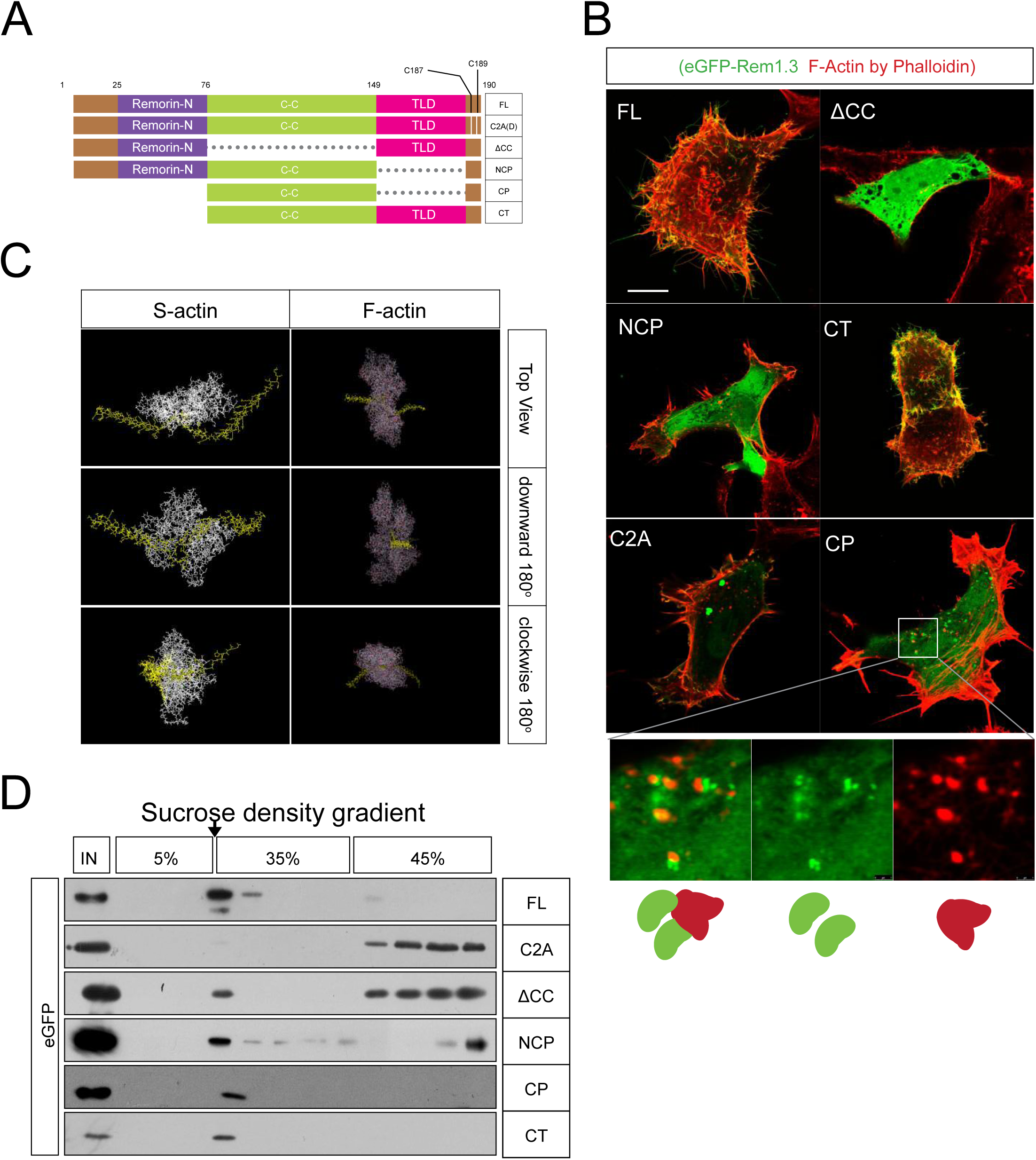
Remorin forms PD-like RFs in mammalian cells. (A) Remorin truncated mutation pattern diagram, the number represents the amino acid position, C represents cysteine. (B) Remorin truncated mutant-transfected in HEK293 cells (green: REM1.3; red: phalloidin-stained F-actin; blue: DNA; scale bar, 10 μm). (C) Predicted structures of REM1.3 protein docking to actin oligomer from PDB (1ATN, 1J6Z, 3EL2 and 4A7N). (D) Western blot analysis of the extracts of lipid microdomains (Arrow indicates) from truncated or mutated eGFP-remorin-transfected HEK-293 cells, a. FL, b. C2A, c. delete 77-149 (ΔCC), d. 1-149 and 186-190 (NCP), e. 77-149 and 186-190 (CP), f. 77-190 (CT). The upper panel indicates the sucrose gradient. IN is an unisolated start sample.

Lack of the N-terminus of REM1.3 (CT) does not obviously affect the remorin localization pattern (Fig 3B and S6). On the other hand, when we truncated most of the N-terminus containing the coiled-coil domain (T) or truncated only four amino acids in the C-terminus (1-186), the distribution pattern of REM1.3 significantly changed into a disperse eGFP signal. Lipid microdomain localization of these two truncated versions was also abolished (Fig 3D and S7), indicating that the coiled-coil domain and the four amino acids in the C-terminus are essential for the formation of RFs. Consistently, a CC domain deletion mutation (ΔCC) could not form RFs (Fig 3A-C).

To study the contribution of the TLD domain (aa 150-186), we linked the four amino acids of the C-terminus directly to the N-terminus with the coiled-coil domain (NCP). This truncation showed a disperse distribution but did not affect the protein lipid microdomain localization. Further deletion of the N-terminal amino acids 1-76 (CP) showed a lipid microdomain distribution but, notably, also small uniform punctate distribution in cells. Phalloidin staining demonstrated that those special structures are adjacent to, but not overlapping with, the F-actin dots (Fig 3B). These results suggest that the TLD domain contributes to the formation of the long RF helix and retains the ability to associate with actin oligomers as an initiative structure of RFs. The observation that the CP version cannot form shaped structures also indicated that the N-terminus 1-76 of REM1.3 is an inhibitory domain for RF formation (Fig 3B).

The lack of coiled-coil domain, mutation of the cysteine at the C-terminal anchor or deletion of the TCHP-like domain (TLD) impairs RF biogenesis. Removing the N-terminal domain from the TCHP-like-domain-lacking REM1.3 can still form the initiatory structure of RFs next to F-actin but results in RFs extending to a lesser degree. This indicates that the coiled-coil domains interact with F-actin and form an RF organization center near the cell membrane, which is responsible for RF initiation. The docking data based on Remorin’s predicted structure and the F-actin crystal structure (http://hexserver.loria.fr/) also support the above notion (Fig 3C). So TCHP-like domain is likely to be responsible for the extension of RFs.

To further study the molecular mechanism of remorin binding to lipid microdomains and their contributions to the formation of RFs, we mutated cysteine 187 or 189 to alanine (Ala, A) or aspartic acid (Asp, D) separately or together. The distribution of REM1.3 changed to inclusion bodies in the cytoplasm (Fig S6), and the proteins disappeared from the lipid microdomains (Fig 3D and S7). This corresponded to our results from the palmitoylation study (Fig S3B). We also linked the four amino acids CGCF of the C-terminus of REM1.3 (aa 187-190) directly into the tail of eGFP; ∼30% eGFP proteins were partially associated with the lipid microdomains (Fig 3D and S7) but showed a dispersed localization (Fig S6), which suggests that those four amino acids are sufficient to recruit the protein onto the lipid microdomains.

To test whether that the knowledge obtained from animal cells is also relevant in plants, we transformed different truncated versions of *REM1.3* driven by a CaMV35S promoter into WT *Arabidopsis*. The results showed that the full-length (FL) protein and the C-terminus (CT) localized at the plasma membrane (PM) without any nuclear signals (Fig S8). In contrast, the CP truncation (aa 77-149+187-190) showed uniform punctate structures (Fig S8), which is consistent to our observations in mammalian cells (Fig S8). Further TEM observation revealed that the FL, CT and CP, led to the dark staining bodies (osmiophilic globules) as observed in the *rem1.2 rem1.3* cells (Fig S1 and S8), which might be due to the ectopic overexpression of *REM1.3*.

### Deep structures of RFs by super-resolution and electron microscopy

To determine the fine structure of plant-remorin induced RF in aminal cells, we use structured-illumination microscopy (SIM), a super-resolution technique evaluated the detailed structure of RFs by eGFP-REM1.3. Refined images (Fig 4A) indicate that the two eGFP-remorin parallel spiral helices twin AFs with an external diameter of 103.34±5.95 nm and a pitch of 140.11±27.21 nm, which is further confirmed by quantification analysis (Table S1). We also used another super-resolution microscopy with a different principle from SIM, stochastic optical reconstruction microscopy (STORM). Both N-Storm (Fig 4B and S4) and C-Storm (Fig 4C) images showed that eGFP-REM1.3-labelled RFs were spiral helices, consistent with the SIM results. These observations revealed striking similarities between RFs and PD structures.

**Fig 4.**
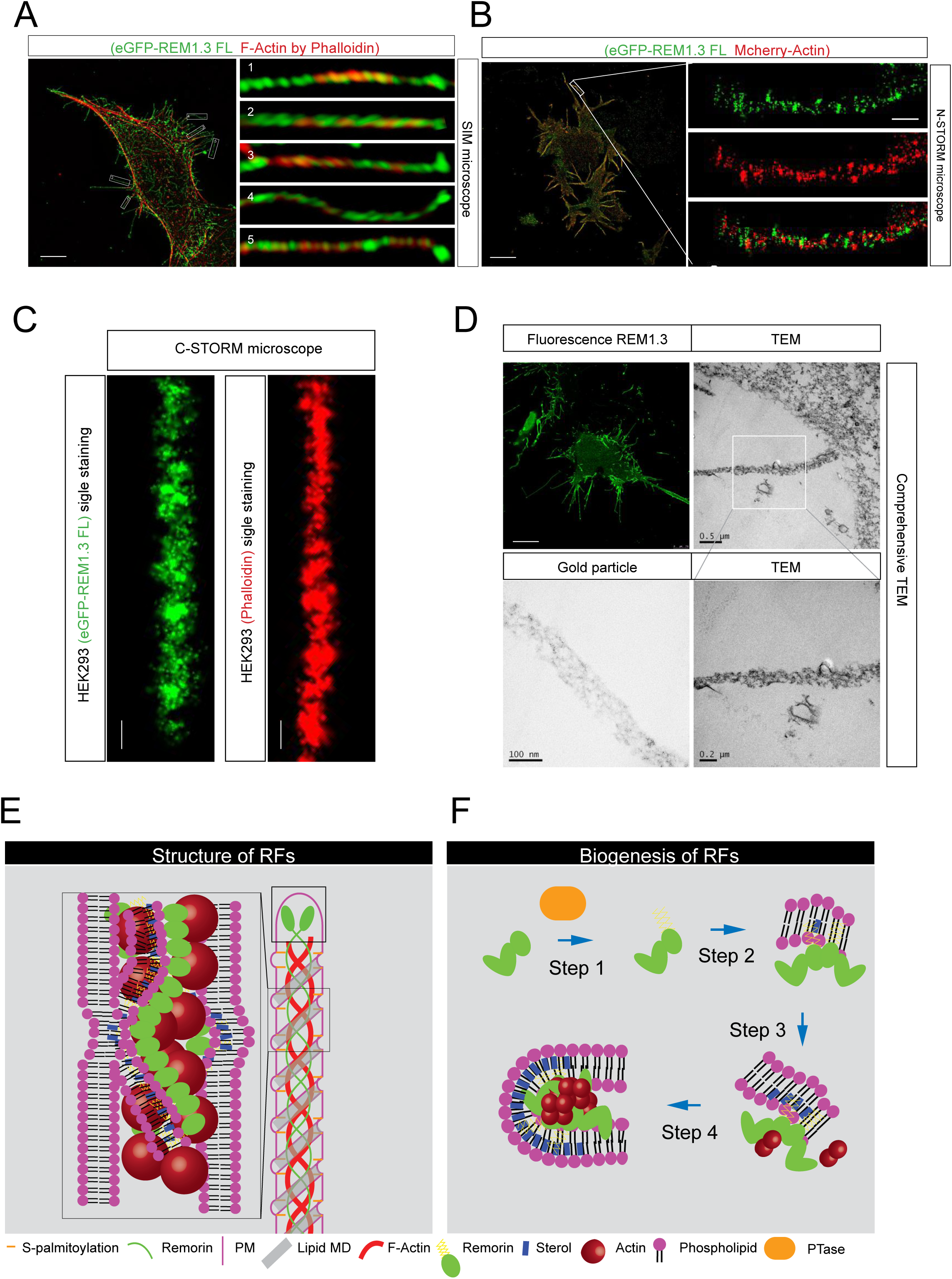
Ultrastructural analysis of RFs. (A) SIM image of eGFP-REM1.3-transfected HEK-293 cell (green: REM1.3; red: phalloidin-stained F-actin; blue: DNA; scale bar, 10 μm). (B) N-STORM images of eGFP-Remorin and Actin-mCherry co-transfected HEK-293 cell stained by activator-reporter paired secondary antibodies (green: 405-647; red: Cy3-647; scale bar, 10 μm). (C) C-STORM images eGFP-Remorin-transfected HEK-293 stained by a GFP antibody and ALEXA-647-labeled secondary antibody (green: REM1.3; red: phalloidin-stained F-actin; scale bar, 100 nm). (D) Comprehensive TEM images of RFs. Fluorescein and nano-gold particle double-labeled secondary antibodies were used to stain RFs in HEK293 cells. Fluorescence signal was collected by confocal microscopy and then analyzed by TEM (green: REM1.3; gray: TEM). (E) A structural cartoon of RFs. RFs are single- or multiple-microfilament structures formed by remorin on the surface of mammalian cells. As seen from the thumbnail on the right, the green remorin and red actin filaments form a double helix. Remorin interacts with the lipid microdomains on the cell membrane (PM) due to its palmitoylation. Therefore, we speculate that the lipid microdomains (Lipid MD) on the surface of the RFs are also helically distributed. The RFs’ head has remorin’s congregation, presumably the center responsible for extending the RFs. On the left is an enlarged fine structure model. (F) A schematic diagram of RFs biogenesis. Remorin modification by palmitoyltransferase is the first step. The second step is remorin and cholesterol-rich cell membrane forming lipid microdomains. In the third step, remorin and lipid microdomains form multimers with actin. In the fourth step, as the RFs polymerize, they begin to extend further to form long RFs.

Moreover, a comprehensive transmission electron microscopy (TEM), which allows us to combine fluorescence imaging and TEM in the same sample by using a fluorescein and nano-gold particle double-labeled secondary antibody, clearly demonstrated that the fluorescence signal of RFs was observed as gold-particle-positive helical structures on TEM (Fig 4D). This further confirmed the spiral structure of RFs determined by super-resolution microscopy.

In summary, these super-resolution and ultrastructure studies revealed that the functional, plant remorin-induced intercellular connections in animal cells show striking ultrastructural similarities to plant plasmodesmata.

## Discussion

### Plant remorin induces functional intercellular connections in animal cells

Previous studies established that plant specific remorins are detected at and functionally important constituents of PD (6, 12, 17). Using a genetic approach, we provided further evidence that remorin regulates the connectivity of the PD confirming remorins as crucial structural and functional components of PD intercellular connections in plants.

Unexpectedly, expressing remorin in animal cell cultures induced functional, PD-like intercellular connections that are clearly distinct from the previously reported well known filamentous cell surface structures, such as cilia, flagella or intermediate filaments (18); therefore we called them RFs.

RFs perserve a high degree of similarity with PD in function. Transporting proteins in the endoplasmic reticulum system is an important functional feature of plant PD. Our experiments show that RFs can not only connect two cardiac muscle cells, and communicate signals between cardiac muscle cell populations and keep them in synchronous rhythm. In addition, RFs can also communicate the biological macromolecules in two animal cells. In our system, RFs enable the provider cells to transport the Cre protein existing in the endoplasmic reticulum to the recipient cells, cut the LoxP motif, and make fluorescence protein expression. An ER-distributed protein is indeed transferred from donor cells to acceptor cells in the presence of remorin, which is very similar to the classic function of PD (1–3).

### Structural similarity between plant PDs and RFs in animals

Here we identified RFs are assembled from remorin, i.e., a quadruple actin-remorin helix (double actin helices × double remorin helices) skeleton for RFs (Fig 4E). Although RFs formed by remorin consists of AFs, it is different from filopodia and tunneling-nanotubes (19). RFs do not have filopodia or tunneling-nanotube marker proteins (3, 19, 20). Moreover, RFs are swollen at both ends, which is different from the classic structure of tunneling-nanotubes, but it is very similar to the structure of a plasmodesma (3). This suggests that a single remorin protein is sufficient for the initiation and maintenance of the core PD structure.

The similar results obtained from both plant and mammalian cells are mutually supportive; for instance, the REM1.3(CP) mutated version exhibited similar behavior in both animal and plant cells linking the cytoskeleton to the cell membrane and stretching it into a filamentous form. Based on these observations, we propose a hypothesized model for PD biogenesis (Fig 4F). Remorin binds to cell membrane through lipid modification, forming a local membrane nanodomain. Meanwhile, remorin also binds the actin filaments, perhaps guiding their direction of extension. This dual function of remorin organizes a local filament, providing the force to remodel the membrane to form a tubular PD structure.

### Evolutionary relation of plant plasmodesmata and animal migrasomes

Migrasome is a newly discovered filamentous structure of animal cells (19, 20). It is related to the cell movement and plays a role in intercellular communication in animal cells. Firstly, migrasome has spherical protrusions on the filaments, and our RFs also have this feature. Second, migrasome has been shown to be encapsulated into cytoplasmic components and released outside the cell (19), and we have observed this phenomenon in RFs, which encapsulates intracellular substances such as DNA and endoplasmic reticulum. Finally, the biochemical properties of RFs are similar to migrasomes, which are rich in Actin and are related to lipid microdomains. Those factors suggest RFs are structurally similar to migrasome (Fig S9). However, biogenesis of migrasome seems to depend on cell movement and division (19, 20), while RFs actively protrude out of cells. Therefore, we hypothesize that remorin may be a factor for multicellular organisms to actively form intercellular connections. This can also be confirmed by the molecular evolution of remorin itself, which firstly appears in multicellular plants above algae, suggesting its causal relationship to multicellular communication. Because animals have evolved a highly developed circulatory system, their permissive capacity between cells is weaker than that of plants. Therefore, the biological mechanism of similar structures is slightly degraded and passive.

## Acknowledgements

This work was supported by National Natural Science Foundation of China [NSFC 31701165, 31370818, 31721001].

## Author contributions

Z. W., S. T. proposed the initial idea of exogenous Remorin in animal cells. Z. W., S. T., K. L and H. X. designed the experiments. Z. W., S. T. and T. L. performed most of the experiments and data analysis. Z. W., T. L. carried out substances transport experiment in animal cells. Z. W., S. T., J.F., K. L and H. X. wrote the manuscript.

## References

1. Brunkard JO, Runkel AM, Zambryski PC (2015) The cytosol must flow: Intercellular transport through plasmodesmata. Curr Opin Cell Biol 35:13–20.

2. Burch-Smith TM, Zambryski PC (2012) Plasmodesmata Paradigm Shift: Regulation from Without Versus Within. Annu Rev Plant Biol 63(1):239–260.

3. Rustom A, Saffrich R, Markovic I, Walther P, Gerdes HH (2004) Nanotubular Highways for Intercellular Organelle Transport. Science 303(5660):1007–1010.

4. Jarsch IK, Ott T (2011) Perspectives on remorin proteins, membrane rafts, and their role during plant-microbe interactions. Mol Plant-Microbe Interact 24(1):7–12.

5. Gui J, Liu C, Shen J, Li L (2014) Grain setting defect1, encoding a remorin protein, affects the grain setting in rice through regulating plasmodesmatal conductance. Plant Physiol 166(3):1463–1478.

6. Raffaele S, Mongrand S, Gamas P, Niebel A, Ott T (2007) Genome-wide annotation of remorins, a plant-specific protein family: Evolutionary and functional perspectives. Plant Physiol 145(3):593–600.

7. Marín M, Thallmair V, Ott T (2012) The intrinsically disordered N-terminal region of AtREM1.3 remorin protein mediates protein-protein interactions. J Biol Chem 287(47):39982–39991.

8. Bariola PA, et al. (2004) Remorins form a novel family of coiled coil-forming oligomeric and filamentous proteins associated with apical, vascular and embryonic tissues in plants. Plant Mol Biol 55(4):579–594.

9. Huang D, et al. (2019) Salicylic acid-mediated plasmodesmal closure via Remorin-dependent lipid organization. Proc Natl Acad Sci U S A 116(42):21274–21284.

10. Gui J, Zheng S, Shen J, Li L (2015) Grain setting defect1 (GSD1) function in rice depends on S-acylation and interacts with actin 1 (OsACT1) at its C-terminal. Front Plant Sci 6(OCTOBER):1–11.

11. Sehnke PC, Chung HJ, Wu K, Ferl RJ (2001) Regulation of starch accumulation by granule-associated plant 14-3-3 proteins. Proc Natl Acad Sci U S A 98(2):765–770.

12. Lynes EM, et al. (2013) Palmitoylation is the switch that assigns calnexin to quality control or ER Ca2+ signaling. J Cell Sci 126(17):3893–3903.

13. Kronberg K, et al. (2007) The silver lining of a viral agent: Increasing seed yield and harvest index in Arabidopsis by ectopic expression of the potato leaf roll virus movement protein. Plant Physiol 145(3):905–918.

14. Chen C, Zhang Y, Zhu L, Yuan M (2010) The actin cytoskeleton is involved in the regulation of the plasmodesmal size exclusion limit. Plant Signal Behav 5(12):1663–1665.

15. Aaziz R, Dinant S, Epel BL (2001) Plasmodesmata and plant cytoskeleton. Trends Plant Sci 6(7):326–330.

16. White RG, Barton DA (2011) The cytoskeleton in plasmodesmata: A role in intercellular transport? J Exp Bot 62(15):5249–5266.

17. Konrad SSA, et al. (2014) S-acylation anchors remorin proteins to the plasma membrane but does not primarily determine their localization in membrane microdomains. New Phytol 203(3):758–769.

18. Konno A, Shiba K, Cai C, Inaba K (2015) Branchial cilia and sperm flagella recruit distinct axonemal components. PLoS One 10(5):1–20.

19. Svitkina TM, et al. (2003) Mechanism of filopodia initiation by reorganization of a dendritic network. J Cell Biol 160(3):409–421.

20. Vignjevic D, et al. (2006) Role of fascin in filopodial protrusion. J Cell Biol 174(6):863–875.

