## Supplemental Material for "Plasmodesmata-like intercellular connections by plant remorin in animal cells"

**Supplementary Materials**

**Materials and Methods:**

*Materials:* Anti-flotillin-1, anti-GFP (mouse monoclonal antibody) antibodies were purchased from Santa Cruz Biotechnology, Inc. (Santa Cruz, CA, USA). The anti-GFP antibody (rabbit monoclonal antibody) was obtained from Cell Signaling Technology. Anti-β-actin, TRITC-phalloidin and HRP-conjugated secondary antibodies were obtained from Sigma (St Louis, MO, USA). Anti-mouse or rabbit fluorescence secondary antibodies and secondary antibodies 405-633 and 561-633 used for N-STORM were purchased or prepared using a kit from Molecular Probe, Invitrogen. Anti-remorin1.3 (AT2G45820) antibody was generated against *Remorin1.3* peptide ESEKSKAENRAQC and KPIEEHTPKKASSGC by Wuxi PharmaTech Corporation Ltd. (Wuxi, China). Fluorescein-10-nm nanogold double-labeled secondary antibody was purchased from Nanoprobes (New York, USA)

*Protein Domain analysis:* REM1.3 was analyzed according to its predicted protein structure on the website InterMap3D or reported domains (Konrad et al., 2014, Perraki et al., 2014, Marin et al., 2012). The predicted 3D-structure of remorin1.3 coiled-coil domain (aa 77-147) is shown in Figure S3B. We Blasted REM1.3 N-terminal domain (aa 1-76), coiled-coil domain (aa 77-147) and C-terminus in mammal protein database in NCBI. The results showed that REM1.3 coiled-coil was homologous with tropomyosin, an actin regulator, which was conserved in eukaryotes(Goodson and Dawson, 2006). Subdivided motifs of coiled-coil using predicted 3D structure contain three obvious domains, helix aa77-90, loop 91-106 and helix 116-143. The C-terminus (aa 55-186) has similarity to a reported keratin filament-binding protein trichoplein (TCHP), which is associated with cilia biogenesis(Ibi et al., 2011); therefore, we name this domain trichoplein TCHP-like domain (TLD). The N-terminal of REM1.3 has no homolog in mammals.

*Plasmid construction: remorin1.3* (*AT2G45820*), *remorin1.2* (*AT3G61260*), *remorin1.4* (*AT5G23750*) and *remorin1.1* (*AT3G48940*) cDNA were cloned from an

*Arabidopsis thaliana* cDNA library and then constructed into a pEGFP-C1 or pmcherry-C1 vector for mammal expression or pHB for plant expression. Truncation, deletion or point mutations were performed by PCR, and the mutated cDNA was tagged by eGFP at the C-terminus. All mutations were confirmed by sequencing.

*Plant materials and growth conditions: Arabidopsis* used in this work were all in the Columbia (Col-0) background. The T-DNA mutants, *rem1.3-1* (SALK_023886C), *rem1.2-1* (SALK_019353C), and *rem1.2-2* (SALK_02445C) were obtained from the *Arabidopsis* Biological Resource Centre(Alonso et al., 2003). *Arabidopsis* seeds were scattered on MS medium plates, stratified for 4 days at 4 °C, and then transferred to 22 °C under long-day (16 h light/ 8 h dark) conditions for growth. Seven-day-old seedlings were then transferred to the soil and grown at the same conditions.

The double knock-out mutant of *rem1.2 rem1.3* was generated by CRISPR-CAS9(Yan et al., 2015). Primers used for genotyping are listed in Table S3.

For transgenic plant construction, versions of REM were fused with a GFP reporter and then sub-cloned into pHB vectors. The resultant plasmids were first introduced into *Agrobacterium GV3101* and then transformed into *Arabidopsis* using the floral dip method(Clough and Bent, 1998).

*Extraction of plant genomic DNA:* A leaf (approximately 50 mg) of adult *Arabidopsis* was kept in a 2-ml EP tube and then treated with 250 μL TPS buffer (1 M KCl, 100 mM Tris-HCl pH = 8.0, 10 mM EDTA) and an iron ball. The samples were ground on a shaker and then incubated at 65 °C for 30 min. After centrifugation at 12000 rpm for 10 min, 200 μl of the supernatant was transferred to a new 1.5 ml EP tube and treated with 200 μl of isopropanol. After mixing, the samples were incubated at room temperature for 20 minutes, centrifuged at 12000 rpm for 10 min and then the supernatant was discarded. The pellet was washed twice with 70% alcohol, dried at 37 °C, and finally dissolved in 50 μl of pure water.

*Arabidopsis starch unloading experiments:* The *Arabidopsis* was cultured for 32 days under normal light conditions for 12 h. Next, the whole plant was covered with aluminum foil, and the light was shielded. After dark treatment for 18 hours, the cut leaves were fixed with absolute ethanol, and the chlorophyll was removed. The leaves were stained with 0.33% Lugol's iodine staining solution (iodine 0.33%, potassium iodide 0.66% diluted in water) for 1 h, washed and photographed (Lee et al., 2011).

*Dye diffusion FRAP experiments: Arabidopsis* thaliana seedlings at 7 days of age were dipped in 1 mM FDA solution for 2 minutes. The stained seedlings used for a FRAP experiment under a laser confocal microscope. The environmental temperature was maintained at 22°C during the entire process.

*Cell culture and transient transfection:* Cells (HEK-293, Cos7) were cultured in Dulbecco's modified Eagle medium (DMEM) supplemented with 10% fetal bovine serum. The transfection was performed with Fugene HD (Promega), following the manufacturer's protocol. The experiments were conducted 48 hours after transfection.

For isolation of primary cardiomyocytes. One-day-old C57BL6 mice were sacrificed. Their hearts were removed, cut in DMEM containing 10 units / ml Collagenase 1 and plated in culture dishes for one hour at 37 °C. Cells with self-paced rhythm were observed within 1 day.

The adenoviral system uses Takara's pAD-easy plasmid system, and the preparation was performed according to the manufacturer's instructions.

*Extraction of lipid microdomains and sucrose density gradient flotation centrifugation:* Transfected cells or smashed were collected in 2 ml ice-cold MBS buffer containing 0.5% Triton X-100, 25 mM MES (pH 6.8), 0.15 M NaCl, 5 mM EDTA, 1 mM PMSF and 2 μl/ml protease inhibitor cocktails 1 and 2. The samples were mixed with an equal volume of 90% sucrose in MBS buffer. The mixture was placed at the bottom of a 12 ml ultra-centrifuge tube and overlaid with 4 ml of 35% sucrose and 4 ml of 5% sucrose in MBS buffer. The gradient was centrifuged at 180,000 ×g for 20 hours in a SW41 rotor (Beckman) at 4 °C, and the fractions were collected after centrifugation(Yao et al., 2009).

*Acyl-Biotin Exchange assay*: Acyl-Biotin Exchange assay was performed according to the method reported by Brigidi and Bamji in 2013(Brigidi and Bamji, 2013). The transfected cells were collected in lysis buffer (50 mM Tris-HCl pH=7.4, 150 mM NaCl, 1% CA-630, 10% glycerol, 1 mM PMSF and 2 μl/ml protease inhibitor cocktails 1 and 2). eGFP-fused remorin was immunoprecipitated by GFP antibodies, and lysis buffer plus 50 mM NEM was added to the beads and incubated for 1 h at 4 °C. After washing the beads with the lysis buffer plus 10 mM NEM and 0.05% SDS, the lysis buffer plus 1 M hydroxylamine (HAM) pH=7.2 were added to beads and incubated at RT for 50 min. The lysis buffer plus a sulfhydryl-reactive biotinylation reagent biotin-BMCC at 2 μM pH=6.2 were added to beads to modify the cysteine residue's thiol groups freed by HAM cleavage. Streptavidin-HRP was used in Western blotting to detect the biotinylated protein.

*Western blotting, confocal microscopy and structured-illumination microscopy:* Western blotting was performed as previously described(Wang et al., 2008). For immunofluorescence imaging, the cells were fixed with 4% paraformaldehyde in phosphate-buffered saline (PBS) and permeated with 0.1% Triton X-100 and 3% bovine serum albumin in PBS. The cells were then incubated with fluorescein-conjugated dyes and photographed using Leica SP8 laser scanning confocal microscope or structured-illumination microscopy (Carl Zeiss AG, Germany). For cells expressing fluorescent-protein-tagged protein, the paraformaldehyde fixed cells were visualized directly by confocal microscope(del Pozo et al., 2005, Li et al., 2009).

*Storm microscopy:* Cultured cells on glass-bottom petri dishes were fixed, permeated and blocked using the immunofluorescence protocol, and the cells were incubated in primary antibodies at RT for 2 h.

For C-Storm imaging, cells were incubated with ALEXA-647-labeled secondary antibody at RT for 1 h, fixed with 4% paraformaldehyde at RT for 10 min, and washed before use. STORM Image Buffer were prepared according to the manual from Nikon and a report from the Zhuang Lab(Shim et al., 2012). Stocking buffer was prepared first: Buffer A containing 10 mM Tris (pH 8.0), 50 mM NaCl; Buffer B containing 50 mM Tris (pH 8.0), 10 mM NaCl, 10% glucose; GLOX solution containing 14 mg glucose oxidase, 50 μl catalase (17 mg/ml) and 200 μl Buffer A; 1 M MEA (1 ml) with 77 mg MEA solute in 1.0 ml 0.25 M HCl. For STORM imaging, the Image Buffer was prepared by adding 70 μl 1 M MEA to 620 μl Buffer B in a 1.5 ml tube. The cells were immersed in Image Buffer, and the STORM signal was detected at 647 nm wave length for approximately 300 frames and analyzed on the Nikon STORM platform.

N-STORM can perform double-channel super-resolution imaging by staining the cells with activator-reporter paired secondary antibodies 405-647 and Cy3-647. When performing N-Storm imaging, 405-nm and 546-nm activated signals were collected circularly. The other steps are the same as for C-Storm.

*Fluorescence-electron microscopy*: Cultured cells on glass-bottom petri dishes were fixed, permeated and blocked following the immunofluorescence protocol. The cells were incubated with primary antibodies and then a fluorescein-10-nm nanogold double-labeled secondary antibody and were photographed by laser scanning confocal microscope. After fluorescence imaging, the cells were post-fixed by 1% osmium tetroxide in PBS buffer for 2 hours at 4 °C. After washing 3 times with distilled water, the samples were dehydrated in ethanol and acetone: 30% ethanol, 50% ethanol and 70% ethanol each for 15 minutes; 80% ethanol and 95% ethanol each for 20 minutes; 100% ethanol twice for 20 minutes each; and 100% acetone three times each for 30 minutes. The dehydrated samples were infiltrated with 1:2 mixture of acetone:Epon 812 ethoxyline for 2 hours and pure Epon 812 ethoxyline overnight and embedded in puro Epon 812 ethoxyline resin. After resin polymerization, the glass bottoms were removed by hydrofluoric acid corrosion. Next, the samples were ultrathin-sectioned (DiATOME knife and Leica ultramicrotome), stained with 2% uranyl acetate and 2% lead citrate and observed under a FEI Tecnai G2 Spirit transmission electron microscope.

*Protein structure prediction and docking:* Full-length Remorin1.3 (AT2G45820) protein sequence was analyzed using the website http://www.cbs.dtu.dk/services/InterMap3D/ with a default parameter setting. The result of the predicted PDB file was visualized in Jmol or submitted as a ligand with receptors of actin 3D structure from PDB (RCSB Protein Data Bank) (1ATN, 1J6Z, 3EL2 and 4A7N) into the protein molecular docking website http://hexserver.loria.fr/ with a default parameter setting(Ghoorah et al., 2013). The docking result was visualized in the 3D Molecule Viewer or analyzed in the WinCoot software.

*Transgenesis of zebrafish:* Fertilized wild-type AB zebrafish eggs were microinjected with 25 ng/mL plasmids in 0.4 M KCl with phenol red solution (Gibco), cultured in E2 media, and observed at 6 h and 24 h after injection.

**Supplementary Figure Legends**

**Fig S1.** Deficiency of remorin leads to PD abnormality in *Arabidopsis*.

A. A schematic diagram of the structures of *REM1.2* and *REM1.3* and the related mutants *rem1.3* and *rem1.2.*

B. Western blotting indicates knock-out of REM1.2 and REM1.3 in *rem1.2*, *rem1.3*, and *rem1.2 rem1.3* mutants using α-REM1.3 / 1.2 antibody, controlled by Coomassie blue staining.

C. The rosette size of *rem1.3. rem1.2* double mutant was smaller than that of WT. Thirty-day-old plants are shown. *rem1.3, rem1.2* and *DKO* (double knock-out).

D. Deficiency of remorin impairs starch transport in *Arabidopsis*. After 45 days of growth of wild-type WT, *rem1.3, rem1.2* and *DKO*, the plants were kept in the dark for 18 h. The leaves were fixed in ethanol and subjected to Logul's iodine staining. The blue-black portion indicates starch accumulation. The images on the right show the leaves without starch accumulation (WT) and starch accumulation (*DKO*) as standards. The bar chart shows the statistical result of starch accumulation (%). Four leaves were randomly selected from one plant, and the study was repeated on 15 plants per line.

E. Dye diffusion experiment for WT and *DKO*, 7-day-old plants were stained with 1 mM FDA solution for 2 minutes. Roots were used for a FRAP experiment under a laser confocal microscope. Heat map shows the fluorescence degree of recovery.

F. Remorin-knockout plants show a decrease of PD and an increase of osmiophilic particles. TEM images of remorin-KO plants. Upper panels are low-magnification images of WT, *rem1.3, rem1.2* and *DKO*; bottom panels are high-magnification images. Green and orange pseudo colors indicate the PD and osmiophilic particles. The bar chart presents the statistical result of the TEM images (PDs and Dots Numbers ± SD). Green and orange bars indicate the number of PD and deep dyeing globules in every two cells, respectively.

**Fig S2.** Remorin can be heterogeneously expressed in mammalian cells with correct localization and modification. A. Western blot results of the extracts of lipid microdomains (indicated by arrow) from eGFP-REM1.3-transfected or mCherry-remorin (REM1.2, 3, 4)-transfected HEK-293 cells and *Arabidopsis* (AT). The upper panel shows the sucrose gradient, and the right panel indicates the antibodies used: flotillin-1 (Flot1) is a lipid microdomain marker (Otto and Nichols, 2011); and actin is a cytosolic protein, which is not localized in lipid microdomain fractions (Liu and Pilch, 2008). B. Detection of palmitoylation of mutated-eGFP-REM1.3-transfected HEK-293 cells using the Acyl-Biotin Exchange assay. C187A, C189A and 2A represent individually mutated cysteine 187 or 189 to alanine or double mutated to alanine. HAM + or – represent addition or no addition of acyl cutting reagent hydroxylamine. Palmitoylation was detected by affiliative blotting with streptavidin-HRP (Biotin), and the PVDF membrane was stripped and subjected to Western blotting for eGFP.

**Fig S3.** RFs and A. B. actin co-localized rather than C. microtubules, D. intermediate filament or E. filopodia. Confocal micrograph of HEK-293 cells transfected with eGFP-Rem1.3 with staining of F-actin (phalloidin) and microtubules by tubulin antibody (tubulin), filopodia or microvilli marker protein Fascin (Fascin), and intermediate filament protein marker Vimentin (Vimentin).

**Fig S4.** N-STORM images of eGFP-Remorin and Actin-mCherry co-transfected HEK-293 cell stained by activator-reporter paired secondary antibodies (green: 405-647; red: Cy3-647; bar = 10 μm). Squares and arrows indicate the axis of rotation and angle.

**Fig S5.** Remorin truncation and point mutation pattern diagram. RFs formation and LR localization represent whether the mutant can form RFs and localize in the lipid microdomains.

**Fig S6.** The function of various REM1.3 mutants in RF biogenesis in eGFP-remorin-transfected HEK-293 cells. Truncated or mutated eGFP-remorins (green) were transformed into HEK-293 cells, and then stained by phalloidin (red). Some results have been shown in the main text figures.

**Fig S7.** Western blot analysis of the extracts of lipid microdomains (Arrow indicates) from truncated or mutated eGFP-remorin-transfected HEK-293 cells, a. FL, b. C187A, c. C189A, d. C187 and 189 to A (C2A), e. 1-186, f. 1-149, g. 77-149 and 186-190 (CP), h. 1-149 and 186-190 (NCP), i. 77-149 (CC), j. delete 77-149 (ΔCC), k. 77-190 (CT), l. 187-190, m. 90-190, n. 1-76 (N), o. 150-190 (T). The upper panel indicates the sucrose gradient. IN is an unisolated start sample. The 5% and 35% interfaces are where the lipid microdomains are located; the right panel indicates the antibodies used: eGFP (eGFP). Some results have been shown in the main text figures.

**Fig S8.** Remorin RF formation domains in mammalian cells related to PD features in plant. Construction of 35S promoter eGFP-remorin-1 transgenic lines in wild-type *Arabidopsis*. The T1 generation plants were grown for 12 days, a laser scanning confocal microscope was used to observe the leaves, bar=20 μm. Location of stomatal guard cell nucleus can be used to indicate that the protein is not located in the cell membrane.

**Fig S9.** Morphological model diagram of PD and animal RFs or migrasome. Migrasome or RFs encapsulate the inner-membrane system or cytoplasm and present an expanded beads morphology. Each expanded bead can be seen as a cell of plant, and the plant has cell wall, so it can be that the cell wall pulls the shape of the expanded bead into a cube as a plant cell morphology. Dotted line boxes indicated that plant cells similar to migrasome.

#### References


ALONSO, J. M., STEPANOVA, A. N., LEISSE, T. J., KIM, C. J., CHEN, H., SHINN, P., STEVENSON, D. K., ZIMMERMAN, J., BARAJAS, P. & CHEUK, R. 2003. Genome-wide insertional mutagenesis of Arabidopsis thaliana. *Science,* 301**,** 653-657.

BRIGIDI, G. S. & BAMJI, S. X. 2013. Detection of protein palmitoylation in cultured hippocampal neurons by immunoprecipitation and acyl-biotin exchange (ABE). *Journal of visualized experiments: JoVE*.

CLOUGH, S. J. & BENT, A. F. 1998. Floral dip: a simplified method forAgrobacterium - mediated transformation ofArabidopsis thaliana. *The plant journal,* 16**,** 735-743.

DEL POZO, M. A., BALASUBRAMANIAN, N., ALDERSON, N. B., KIOSSES, W. B., GRANDE-GARC A, A., ANDERSON, R. G. & SCHWARTZ, M. A. 2005. Phospho-caveolin-1 mediates integrin-regulated membrane domain internalization. *Nature cell biology,* 7**,** 901.

GHOORAH, A. W., DEVIGNES, M.-D., SMAIL-TABBONE, M. & RITCHIE, D. W. 2013. Protein docking using case-based reasoning. *Proteins-Structure Function and Bioinformatics,* 81**,** 2150-2158.

GOODSON, H. V. & DAWSON, S. C. 2006. Multiplying myosins. *Proceedings of the National Academy of Sciences of the United States of America,* 103**,** 3498-3499.

IBI, M., ZOU, P., INOKO, A., SHIROMIZU, T., MATSUYAMA, M., HAYASHI, Y., ENOMOTO, M., MORI, D., HIROTSUNE, S. & KIYONO, T. 2011. Trichoplein controls microtubule anchoring at the centrosome by binding to Odf2 and ninein. *J Cell Sci,* 124**,** 857-864.

KONRAD, S. S., POPP, C., STRATIL, T. F., JARSCH, I. K., THALLMAIR, V., FOLGMANN, J., MAR N, M. & OTT, T. 2014. S - acylation anchors remorin proteins to the plasma membrane but does not primarily determine their localization in membrane microdomains. *New Phytologist,* 203**,** 758-769.

LEE, J.-Y., WANG, X., CUI, W., SAGER, R., MODLA, S., CZYMMEK, K., ZYBALIOV, B., VAN WIJK, K., ZHANG, C. & LU, H. 2011. A plasmodesmata-localized protein mediates crosstalk between cell-to-cell communication and innate immunity in Arabidopsis. *The Plant Cell,* 23**,** 3353-3373.

LI, K., YAO, W., ZHENG, X. & LIAO, K. 2009. Berberine promotes the development of atherosclerosis and foam cell formation by inducing scavenger receptor A expression in macrophage. *Cell research,* 19**,** 1006.

LIU, L. & PILCH, P. F. 2008. A critical role of cavin (polymerase I and transcript release factor) in caveolae formation and organization. *Journal of Biological Chemistry,* 283**,** 4314-4322.

MARIN, M., THALLMAIR, V. & OTT, T. 2012. The Intrinsically Disordered N-terminal Region of AtREM1.3 Remorin Protein Mediates Protein-Protein Interactions. *Journal of Biological Chemistry,* 287.

OTTO, G. P. & NICHOLS, B. J. 2011. The roles of flotillin microdomains–endocytosis and beyond. *J Cell Sci,* 124**,** 3933-3940.

PERRAKI, A., BINAGHI, M., MECCHIA, M. A., GRONNIER, J., GERMAN-RETANA, S., MONGRAND, S., BAYER, E., ZELADA, A. M. & GERMAIN, V. 2014. StRemorin1. 3 hampers Potato virus X TGBp1 ability to increase plasmodesmata permeability, but does not interfere with its silencing suppressor activity. *FEBS letters,* 588**,** 1699-1705.

SHIM, S.-H., XIA, C., ZHONG, G., BABCOCK, H. P., VAUGHAN, J. C., HUANG, B., WANG, X., XU, C., BI, G.-Q. & ZHUANG, X. 2012. Super-resolution fluorescence imaging of organelles in live cells with photoswitchable membrane probes. *Proceedings of the National Academy of Sciences,* 109**,** 13978-13983.

WANG, W., CHEN, L., DING, Y., JIN, J. & LIAO, K. 2008. Centrosome separation driven by actin-microfilaments during mitosis is mediated by centrosome-associated tyrosine-phosphorylated cortactin. *Journal of cell science,* 121**,** 1334-1343.

YAN, L., WEI, S., WU, Y., HU, R., LI, H., YANG, W. & XIE, Q. 2015. High-Efficiency Genome Editing in Arabidopsis Using YAO Promoter-Driven CRISPR/Cas9 System. *Molecular Plant,* 8**,** 1820-1823.

YAO, Y., HONG, S., ZHOU, H., YUAN, T., ZENG, R. & LIAO, K. 2009. The differential protein and lipid compositions of noncaveolar lipid microdomains and caveolae. *Cell research,* 19**,** 497.

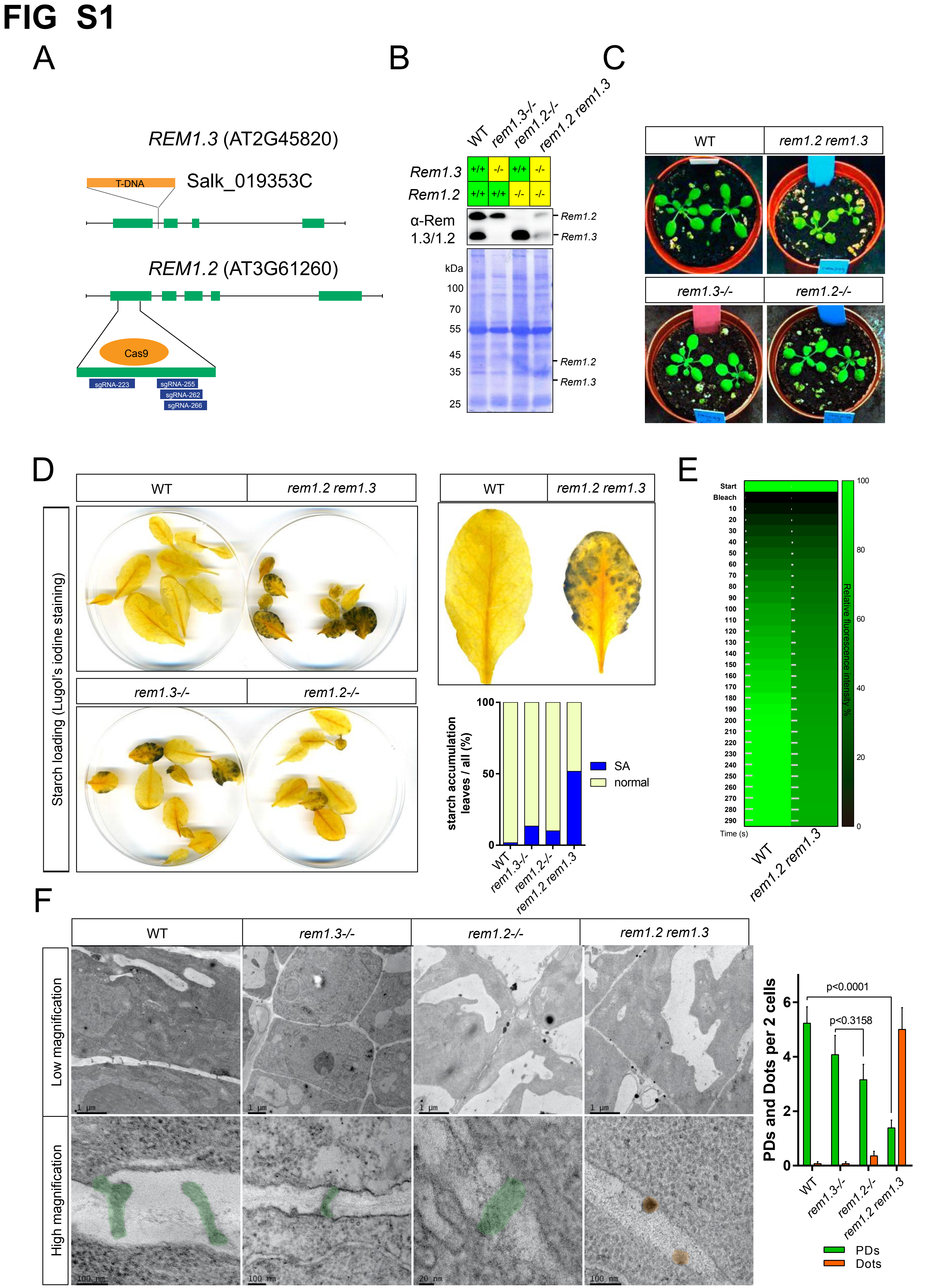
FIG S1
A
REM1.3 (AT2G45820)
T-DNA
Salk_019353C
REM1.2 (AT3G61260)
Cas9
sgRNA-223
sgRNA-255
sgRNA-262
sgRNA-266
B
WT
rem1.3-/-
rem1.2-/-
rem1.2 rem1.3
Rem1.3
Rem1.2
α-Rem 1.3/1.2
kDa
100
70
55
45
35
25
C
D
Starch loading (Lugol's iodine staining)
starch accumulation leaves / all (%)
SA
normal
E
Start
Bleach
Time (s)
Relative fluorescence intensity %
F
Low magnification
High magnification
PDs and Dots per 2 cells
p<0.0001
p<0.3158
PDs
Dots

### FIG S2

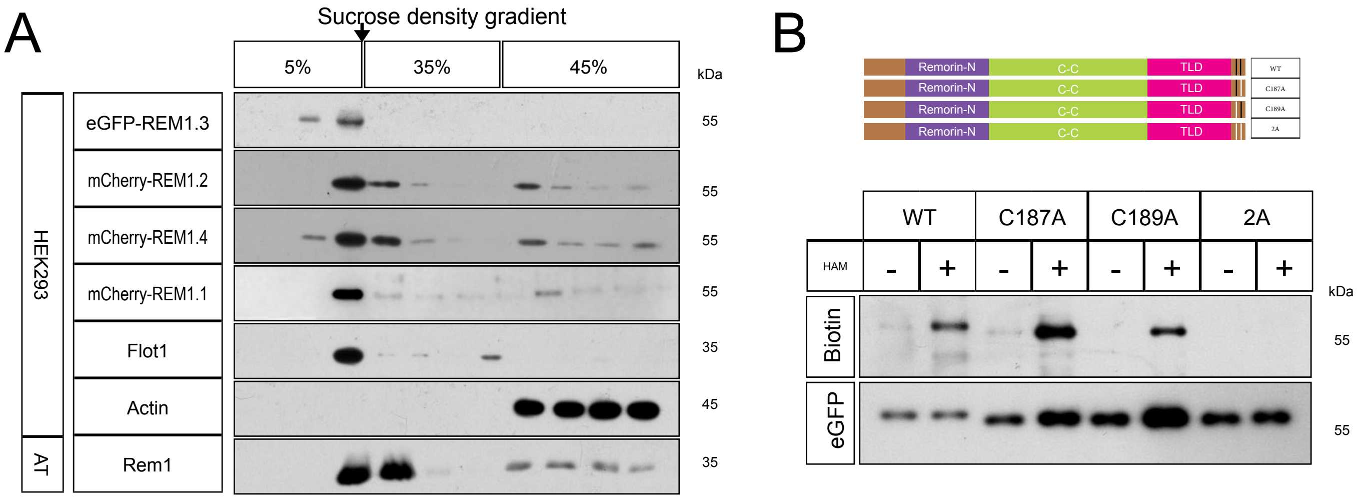
A
Sucrose density gradient
5%
35%
45%
kDa
HEK293
AT
eGFP-REM1.3
mCherry-REM1.2
mCherry-REM1.4
mCherry-REM1.1
Flot1
Actin
Rem1
55
55
55
55
35
45
35
B
Remorin-N
C-C
TLD
WT
C187A
C189A
2A
HAM
-
+
Biotin
eGFP
kDa
55
55

**FIG S3**

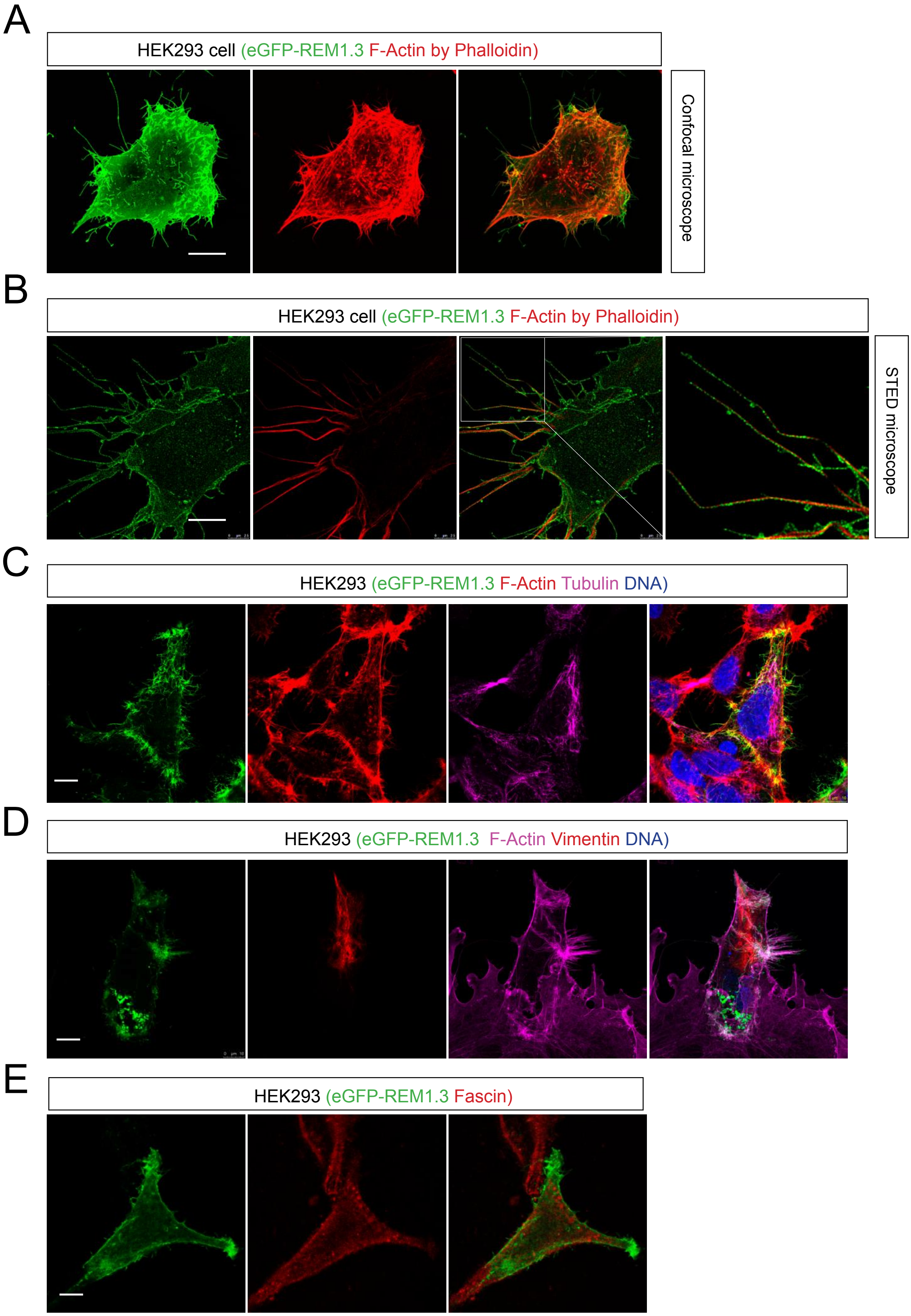

A
HEK293 cell (eGFP-REM1.3 F-Actin by Phalloidin)
Confocal microscope
B
HEK293 cell (eGFP-REM1.3 F-Actin by Phalloidin)
STED microscope
C
HEK293 (eGFP-REM1.3 F-Actin Tubulin DNA)
D
HEK293 (eGFP-REM1.3 F-Actin Vimentin DNA)
E
HEK293 (eGFP-REM1.3 Fascin)

### FIG S4

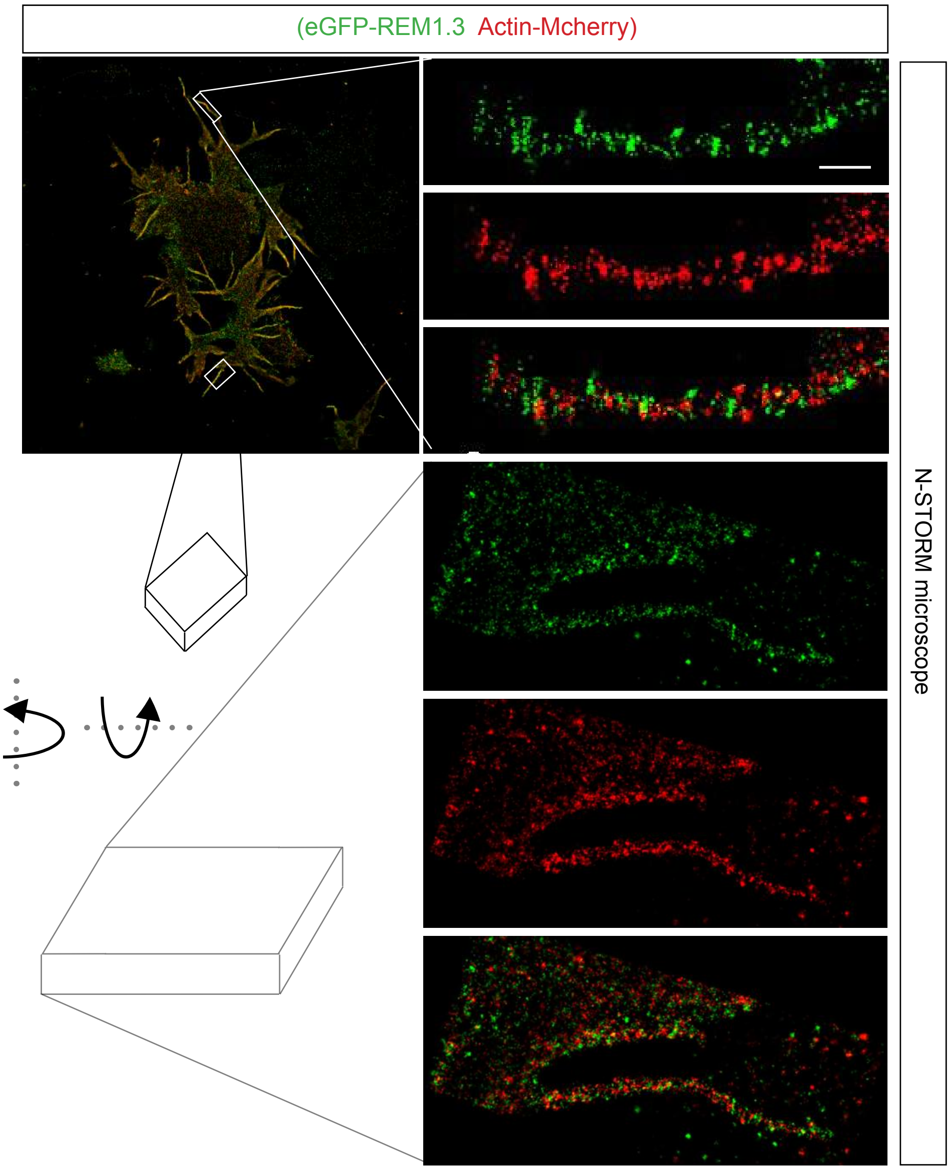

### FIG S5

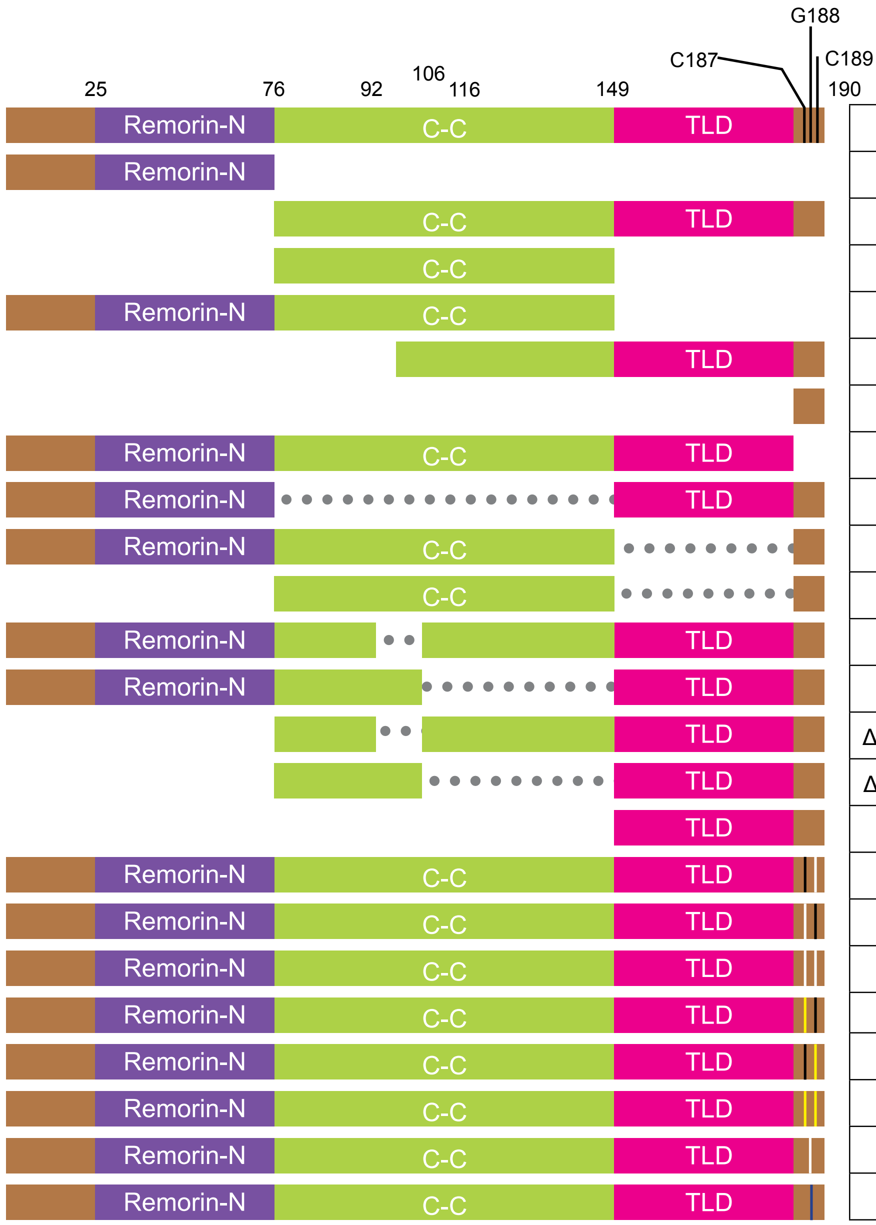


| | RFs formation | LR localization |
|---|---|---|
| FL | Yes | Yes |
| N | No | No |
| CT | Yes | Yes |
| CC | No | No |
| NC | No | No |
| 90-190 | No | Yes |
| 187-190 | No | Partly |
| 1-186 | No | No |
| ΔCC | No | Partly |
| NCP | No | Partly |
| CP | No but special | Yes |
| ΔCC-loop | No | Partly |
| Δ116-147 | No | Partly |
| ΔN+ΔCC-loop | No | Partly |
| ΔN+Δ116-147 | No | Partly |
| T | No | No |
| C187A | Partly | No |
| C189A | Partly | No |
| C2A | No | No |
| C187D | Partly | No |
| C189D | No | No |
| 2D | No | No |
| G188A | Yes | Yes |
| G188Y | Yes | Yes |

### FIG S6

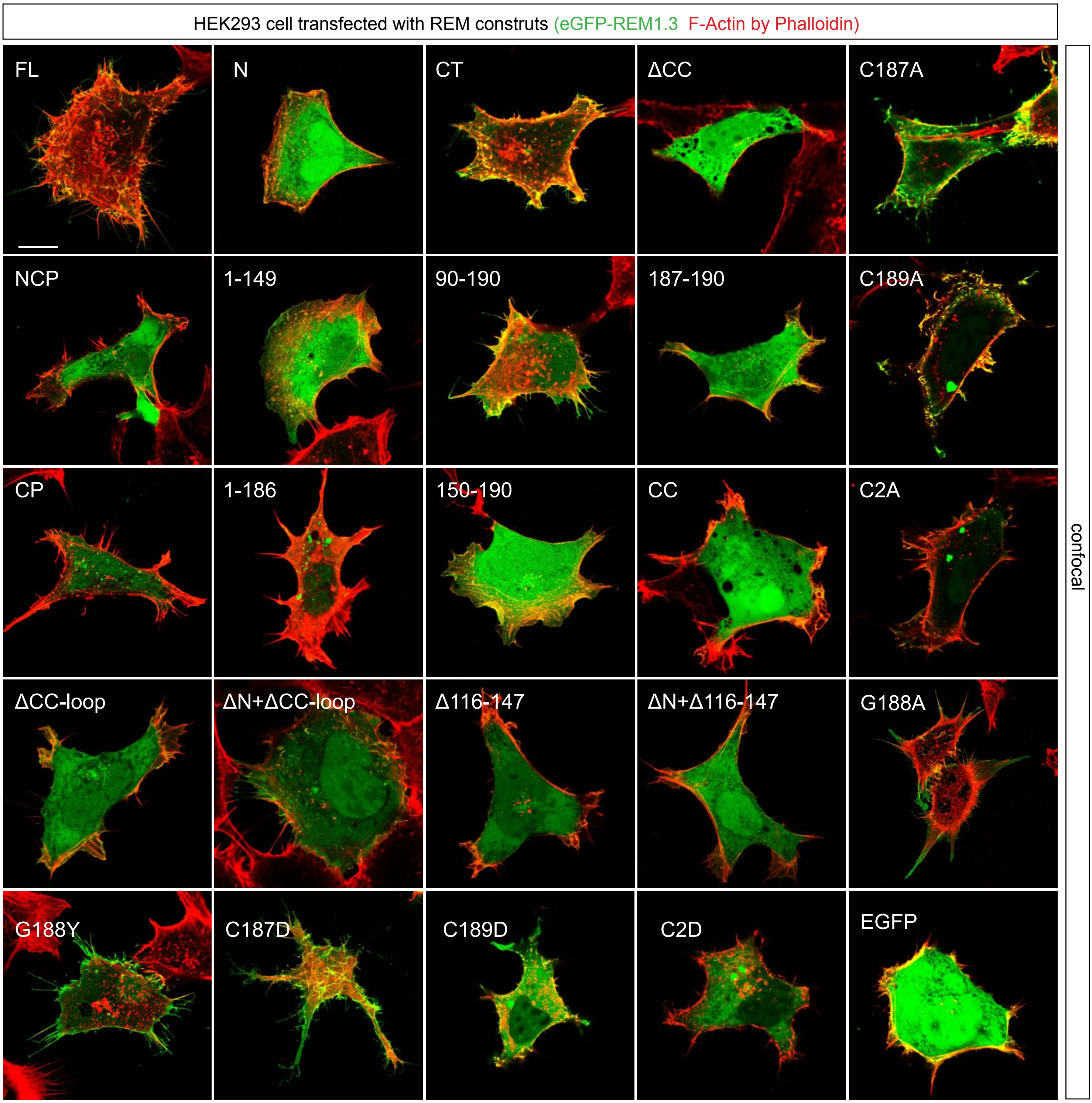
HEK293 cell transfected with REM construts (eGFP-REM1.3 F-Actin by Phalloidin)
FL
N
CT
ΔCC
C187A
NCP
1-149
90-190
187-190
C189A
CP
1-186
150-190
CC
C2A
ΔCC-loop
ΔN+ΔCC-loop
Δ116-147
ΔN+Δ116-147
G188A
G188Y
C187D
C189D
C2D
EGFP
confocal

**FIG S7**

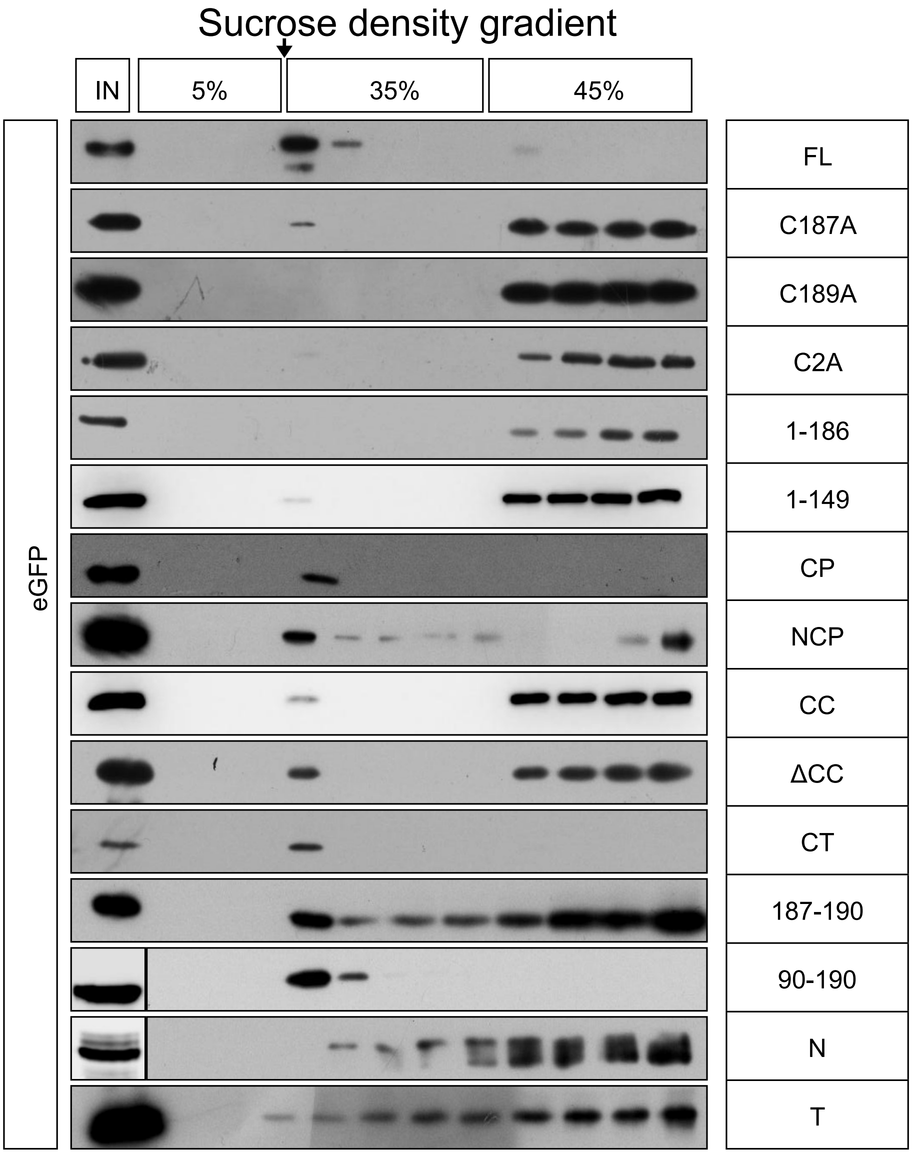

Sucrose density gradient
IN
5%
35%
45%
eGFP
FL
C187A
C189A
C2A
1-186
1-149
CP
NCP
CC
ΔCC
CT
187-190
90-190
N
T

FIG S8

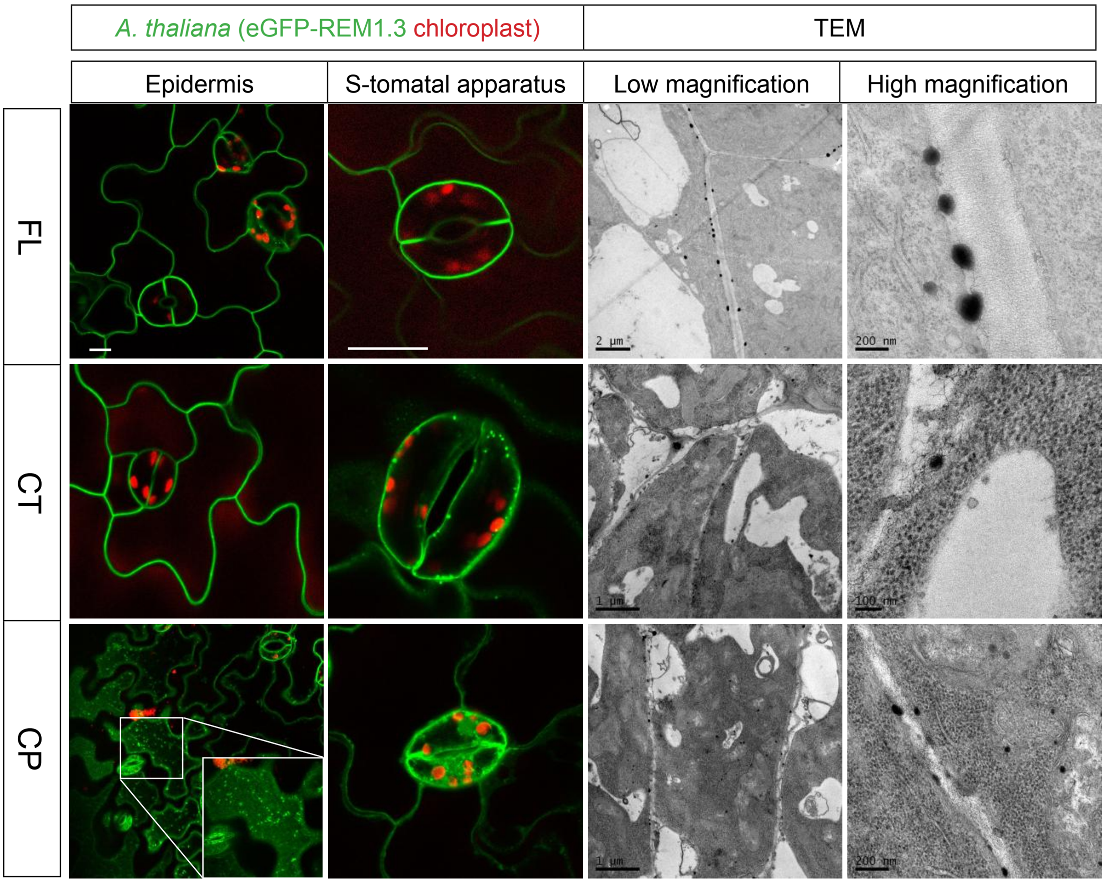

A. thaliana (eGFP-REM1.3 chloroplast)
TEM
Epidermis
S-tomatal apparatus
Low magnification
High magnification
FL
CT
CP
2 µm
200 nm
1 µm
100 nm
1 µm
200 nm

### FIG S9

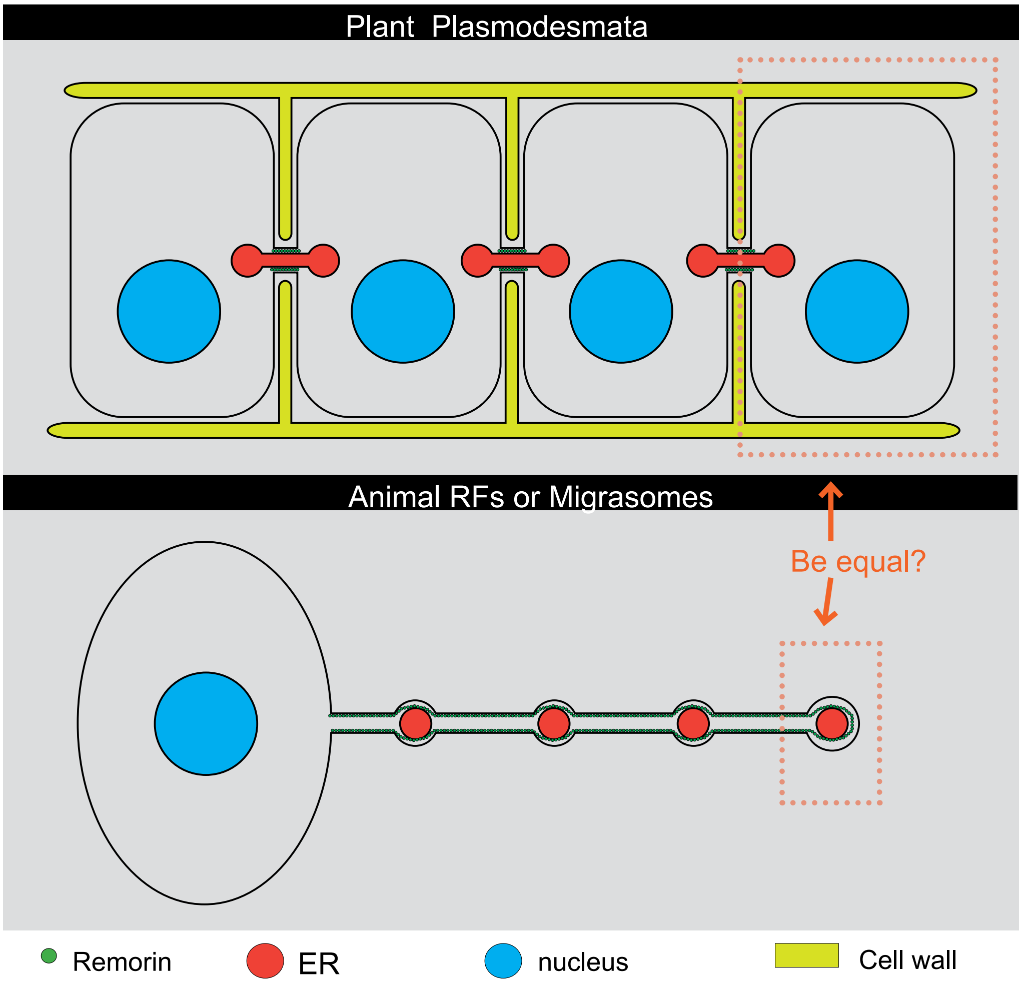
Plant Plasmodesmata
Animal RFs or Migrasomes
Be equal?
Remorin
ER
nucleus
Cell wall

**Table S1. RFs morphology index.**

| **Index** | **Mean** ± SD |
|---|---|
| branch | 1.29±0.31 |
| Length (μm) | 11.54±1.44 |
| external diameter (nm) | 103.34±5.95 |
| inner diameter (nm) | 46.90±3.01 |
| pitch (nm) | 140.11±27.21 |
| lead angle | 73.86°±12.64 |

**Table S2. Domain's contribution for RFs formation of Remorin 1.3**

| Domain | Amino acid | Functions |
|---|---|---|
| N-terminal (N) | 1-76 | Negative |
| Coiled-Coil (CC) | 77-147 | Actin regulation |
| TCHP-like domain (TLD) | 150-186 | Elongation RFs |
| Palmitoylated Tail (P) | 187-190 | LR localization |

**Table S3. Primers used in this study.** Added restriction enzymes underlined.

| Name | Sequence (5'-3') | Enzyme | Plasmid | Usage |
|---|---|---|---|---|
| pEGFPC1-REM1.3-F | cgcggatccCGGTCGCCACCATGGTGAGC | BamHI | pEGFP-C1 | Animal expression |
| pEGFPC1-REM1.3-R | ccgctcgagGATACAAATTCAATAAAGGC | XhoI | pEGFP-C1 | Animal expression |
| bgl2-mcherryc1-REM2-AT3G61260.1-f | GAAGATCTATGGCGGAGGAACAGAAGATAG | BglII | pmcherry-C1 | Animal expression |
| ecor1-mcherryc1-REM2-AT3G61260.1-r | gcggaattcTTAGAAACATCCACAAGTTGCC | EcoRI | pmcherry-C1 | Animal expression |
| bgl2-mcherryc1-REM3-AT5G23750.1-f | GAAGATCTATGGCTGAAGAGGAACCGAAGAAGGT | BglII | pmcherry-C1 | Animal expression |
| ecor1-mcherryc1-REM3-AT5G23750.1-r | gcggaattcTCACATGCATCCGAAAAGCTTTTTGGGA | EcoRI | pmcherry-C1 | Animal expression |
| bgl2-mcherryc1-REM4-AT3G48940.1-f | GAAGATCTATGACTTTAGAGGAGCAGAAGAAAGT | BglII | pmcherry-C1 | Animal expression |
| ecor1-mcherryc1-REM4-AT3G48940.1-r | gcggaattcTCAGAAGAATCCAAATAGCTTGGTTGGA | EcoRI | pmcherry-C1 | Animal expression |
| tag2b-BamH1-f-REM-1 | cgcggatccATGGCGGAGGAGCAAAAGACGAGT | BamHI | pCMV-tag2B | Animal expression |
| GFPC1-Xho1-f-REM-1 | ccgctcgagctATGGCGGAGGAGCAAAAGACGAGT | XhoI | pEGFP-C1 | Animal expression |
| GFPC1-bamH1-r-REM-190 | cgcggatccAGGCTTAGAAACATCCACACGTTG | BamHI | pEGFP-C1 | Animal expression |
| GFPC1-bamH1-r-REM-76 | cgcggatccCAAGTCGGCAAGTATCACATCTC | BamHI | pEGFP-C1 | Animal expression |
| GFPC1-Xho1-f-REM-77 | ccgctcgagctGAAAAAGAGAAGAAAACGTCATTCAT | XhoI | pEGFP-C1 | Animal expression |
| GFPC1-Xho1-f-REM-55 | ccgctcgagctGAGCATACACCTAAGAAAGCTTCAT | XhoI | pEGFP-C1 | Animal expression |
| GFPC1-bamH1-r-REM-149 | cgcggatccCTTGTGGATTGCAGCTACTTTG | BamHI | pEGFP-C1 | Animal expression |
| GFPC1-Xho1-f-REM-150 | ccgctcgagctTTAGCAGAAGAGAAGAGAGCAAT | XhoI | pEGFP-C1 | Animal expression |

| | | | | |
|---|---|---|---|---|
| GFPC1-Xho1-f-REM-90 | ccgctcgagctGAGAGTGAGAAGTCAAAGGCT | XhoI | pEGFP-C1 | Animal expression |
| GFPC1-Xho1-f-REM-100 | ccgctcgagctGCACAAAAGAAGATCTCTGAT | XhoI | pEGFP-C1 | Animal expression |
| GFPC1-Xho1-f-REM-110 | ccgctcgagctTGGGAAAATAGCAAGAAAGCAG | XhoI | pEGFP-C1 | Animal expression |
| GFPC1-Xho1-f-REM-120 | ccgctcgagctGCTCAACTTAGGAAGATCGAGG | XhoI | pEGFP-C1 | Animal expression |
| GFPC1-Xho1-f-REM-130 | ccgctcgagctGAGAAGAAAAAAGCGCAGTAC | XhoI | pEGFP-C1 | Animal expression |
| GFPC1-bamH1-r-REM-186 stop | cgcggatccTTACGTTGCCTTTGGTACTACACCAG | BamHI | pEGFP-C1 | Animal expression |
| GFPC1-bamH1-r-REM-186 | cgcggatccCGTTGCCTTTGGTACTACACCAG | BamHI | pEGFP-C1 | Animal expression |
| C1-bamH1-r-REM-187and189CtoA | cgcggatccTTAGAATGCTCCTGCCGTTGCCTTTGGT | BamHI | pEGFP-C1 | Animal expression |
| C1-bamH1-r-REM-187CtoA | cgcggatccTTAGAAACATCCTGCCGTTGCCTTTGGT | BamHI | pEGFP-C1 | Animal expression |
| C1-bamH1-r-REM-189CtoA | cgcggatccTTAGAATGCTCCACACGTTGCCTTTGGT | BamHI | pEGFP-C1 | Animal expression |
| C1-bamH1-r-REM-149+186-190 | cgcggatccTTAGAAACATCCACACGTCTTGTGGATTGCAGCTACTTTG | BamHI | pEGFP-C1 | Animal expression |
| C1-bamH1-r-REM-187and189CtoD | cgcggatccTTAGAAGTCTCCGTCCGTTGCCTTTGGT | BamHI | pEGFP-C1 | Animal expression |
| C1-bamH1-r-REM-187CtoD | cgcggatccTTAGAAACATCCGTCCGTTGCCTTTGGT | BamHI | pEGFP-C1 | Animal expression |
| C1-bamH1-r-REM-189CtoD | cgcggatccTTAGAAGTCTCCACACGTTGCCTTTGGT | BamHI | pEGFP-C1 | Animal expression |
| REM-D92-147-Rs | AAAGCATGGGAAGAGAGTCACAAGTTAGCAGAAGAG | | pEGFP-C1 | deletion mutation process primers |
| REM-D92-147-Fa | CTCTTCTGCTAACTTGTGACTCTCTTCCCATGCTTT | | pEGFP-C1 | deletion mutation process primers |
| C1-bamH1-r-REM-188GtoA | cgcggatccTTAGAAACAGGCACACGTTGCCTT | BamHI | pEGFP-C1 | Animal expression |
| C1-bamH1-r-REM-188GtoY | cgcggatccTTAGAAACAGTAACACGTTGCCTT | BamHI | pEGFP-C1 | Animal expression |
| REM1-CC187-190-s | gatctTGTGGATGTTTCTAAg | BglII | pEGFP-C1 | deletion mutation process primers |

| | | | | |
|---|---|---|---|---|
| REM1-CC187-190-r | aattcTTAGAAACATCCACAa | EcoRI | pEGFP-C1 | deletion mutation process primers |
| REM1-AA187-190-s | gatctGCAGGAGCATTCTAAg | BglII | pEGFP-C1 | deletion mutation process primers |
| REM1-AA187-190-r | aattcTTAGAATGCTCCTGCa | EcoRI | pEGFP-C1 | deletion mutation process primers |
| REM-D94-103-Fa | GGAAGAGAGTGAGAAGATCTCTGATGTGCATG | | pEGFP-C1 | deletion mutation process primers |
| REM-D94-103-Rs | CATGCACATCAGAGATCTTCTCACTCTCTTCC | | pEGFP-C1 | deletion mutation process primers |
| REM-D116-143-Fa | TGTGGATTGCAGCTACTTTCTTGCTATTTTCC | | pEGFP-C1 | deletion mutation process primers |
| REM-D116-143-Rs | GGAAAATAGCAAGAAAGTAGCTGCAATCCACA | | pEGFP-C1 | deletion mutation process primers |
| EGFPc1topHB-Pst1-f | GCTGCAGCTTTAGTGAACCGTCAGATCCGCT | PstI | pHB | plant expression |
| EGFPc1topHB-Sac1-r | CGAGCTCGCAAATGTGGTATGGCTGATTAT | SacI | pHB | plant expression |
| bgl2-loxp-mcherry-F | GAAGATCTATAACTTCGTATAGCATACATTATACGAAGTTATTCAGATCCGCTAGCGCTACC | BglII | pLVX-IRES-puro | lmlg construction process primers |
| kpn1-mcherry-stop-loxp-R | GGGGTACCATAACTTCGTATAATGTATGCTATACGAAGTTATTCGAGCGGCCTTATTCAGCTAGCTctaCTTGTACAGCTCGTCCATGC | KpnI | pLVX-IRES-puro | lmlg construction process primers |
| ecor1-lmlg-f | CCGGAATTCgcgctaccggactcagatct | EcoRI | pLVX-IRES-puro | lmlg construction process primers |
| xba1-lmlg-r | GCTCTAGActagagtcgcggccgcttta | XbaI | pLVX-IRES-puro | lmlg construction process primers |
| fama(1-40)-cre-Pds | CCACAGCGTCCATCCGCCATGGCACCCAAGAAGAAG | | pLVX-IRES-puro | ERs -Cre construction process primers |
| fama(1-40)-cre-Nua | CTTCTTCTTGGGTGCCATGGCGGATGGACGCTGTGG | | pLVX-IRES-puro | ERs-Cre construction process primers |
| bgl2-Cre-end-r | GAAGATCTGCCACCCTAATCGCCATCTTGCAGCA | BglII | pLVX-IRES-puro | ERs -Cre construction process primers |
| ecoR1-hfam198a-1-f | CCGGAATTCatggcgtcttggctccgg | EcoRI | pLVX-IRES-puro | ERs -Cre construction process primers |
| Not1-hfam198a-1-f | ATAAGAATGCGGCCGCatggcgtcttggctccgg | NotI | pLVX-IRES-puro | ERs -Cre construction process |

| | | | | |
|---|---|---|---|---|
| | | | | primers |
| pYAO-c9-sgRNA-223-s | attgTGTCACGAAAGACGTTGCAG | BsaI | pYao-cas9 | Cas9 for REM1.2 |
| pYAO-c9-sgRNA-223-r | aaacCTGCAACGTCTTTCGTGACA | BsaI | pYao-cas9 | Cas9 for REM1.2 |
| ATrem2-7-26-F | AATATCTTCAATTTTTCATA | | | genome detection for REM1.2 |
| ATrem2-458-477-R | ATTTCAATGACAAATTCTAT | | | genome detection for REM1.2 |
| rem1.3-1_LP | TGATGAATGACGTTTTCTTCTC | | | Genotyping for SALK_023886C |
| rem1.3-1_RP | TTCATAATCCACCTCCCGTCG | | | Genotyping for SALK_023886C |
| LBa1 | TGGTTCACGTAGTGGGCCATCG | | | For SALK mutants |
| LB1 | GCCTTTTCAGAAATGGATAAATAGCCTTGCTTCC | | | For SALK mutants |
| REM1.3_F | TGATGAATGACGTTTTCTTCTC | | | qRT-PCR |
| REM1.3_R | TTCATAATCCACCTCCCGTCG | | | qRT-PCR |
| ACTIN7-F | CCGGTATTGTGCTCGATTCTG | | | qRT-PCR |
| ACTIN7-R | TTCCCGTTCTGCGGTAGTGG | | | qRT-PCR |
